# Efficient detection and typing of phage-plasmids

**DOI:** 10.1101/2025.08.29.673033

**Authors:** Karina Ilchenko, Remy A. Bonnin, Eduardo PC Rocha, Eugen Pfeifer

## Abstract

Phage-plasmids are temperate phages that replicate as plasmids during lysogeny. Despite their high diversity, they carry genes similar to phages and plasmids. This leads to gene exchanges, and to the formation of hybrid or defective elements, which limits accurate detection of phage-plasmids.

To address this challenge, we present tyPPing, an easy-to-use method that efficiently detects and types phage-plasmids with high accuracy. It searches for distinct frequencies and sets of conserved proteins to separate phage-plasmids from plasmids and phages. tyPPing’s strength comes from both its precise predictions and its ability to systematically type phage-plasmids, including the assignment of confidence levels. We tested tyPPing on current databases and a collection of incomplete (draft) genomes. While predictions rely on the quality of assemblies, we detected high-quality phage-plasmids and experimentally proved them to be functional. Compared to other classification methods, tyPPing is designed to detect distinct types of phage-plasmids, and surpasses other tools in terms of sensitivity and scalability. Phage-plasmids are highly diverse, making the systematic identification of new types a difficult task. By combining tyPPing with other tools, however, we show a valuable foundation for addressing this challenge.

## Introduction

Phage-plasmids (or prophage-plasmids, short P-Ps) challenges the traditional dichotomy between a bacteriophage (or phage), which is infectious and often lytic, and a plasmid, which is a stable, autonomously-replicating genetic element. P-Ps are defined as temperate dsDNA phages that typically lack integrases and replicate episomally in bacterial hosts as plasmids. Contrary to integrative temperate phages, the episomal state allows P-Ps to exist as multiple copy elements (per cell). This polyploidy has been reported to promote a gradual spread of mutations, which helps avoid their complete loss through segregational drift (1). As plasmids, P-Ps rely on partitioning systems to ensure their proper segregation to daughter cells (during cell division), and they may encode addiction modules (toxin/antitoxin systems (2, 3)), facilitating genetic stability. Signals such as DNA damage or quorum sensing molecules have been described (4–6) to cause the switch to the lytic cycle, ultimately initiating the production of new progeny and lysis of the host. Most of our current understanding of P-Ps stems from studies on two well-characterized phages, P1 (7) and N15 (8). However, in recent years, a growing body of research has begun to shed new light on their biology and broader impact (9–12).

Separating P-Ps from other phages and plasmids is challenging given their dual nature. For clarity, we will refer to them as P-Ps, and differentiate them from other phages and other plasmids (which will be just called phages and plasmids). P-Ps are reported to be diverse and widespread across bacteria (9). They encode a variety of intriguing traits, such as virulence factors (9, 13) and antibiotic resistance genes (ARGs) (14), with the latter being rare in phages (14, 15). Yet, clinical strains often harbor ARGs that are encoded on P-Ps, suggesting them to have a substantial role in their dissemination (14, 16, 17). P-Ps have homologs in plasmids, which facilitates recombination, and potentially the acquisition of ARGs. They can spread ARGs via lysogenic conversion and remain infective as phages (once arrived in a new host) (14).

We therefore postulate that P-P detection is crucial to understand ARG transmission. Furthermore, as P-Ps possess also homologs in phages, they drive a gene flow between phages, plasmids and P-Ps (18), positioning them as a key exchange point between MGEs. P-Ps exhibit evolutionary plasticity and may evolve to plasmids and to integrative prophages (18). These dynamics suggest a continuous nature of P-Ps, phages and plasmids, and underscore substantial challenges in the precise classification of P-Ps.

No specific tool has been developed to effectively detect P-Ps, with most current methods focusing on the detection of integrative prophages, plasmids, and other MGEs (19–24). We noted that P-Ps are often miss-annotated, solely as phages or plasmids. To tackle this, we screened plasmid and phage databases for P-Ps and created a first P-P catalogue to address their diversity (9). In our approach we annotated plasmid sequences with phage profiles and random forest models, and screened phages for plasmid functions. We complemented our search with an exhaustive literature review, and added cases that were missed by the two approaches. By using the gene repertoire relatedness of P-Ps, we grouped them into well-related types: AB, P1, N15, and five SSU5-related groups. Most of these (91.7%) are predicted to infect Enterobacterales species. The other P-Ps were either singletons (unique P-Ps that are not related to others) or clustered into sparsely-related communities with members that are frequently detected (approx. 1 out of 2) in monoderm species (*Actinomycetia*, *Bacilli*, and *Clostridia*) (9). We found that the P-P types have unique, intriguing characteristics related to their gene repertoires (core and accessory), genomic sizes, and evolutionary histories. Yet, our first approach, despite detecting many P-Ps, faces important drawbacks. First, the detection requires time-consuming and labor-intensive (semi-automated) screens that are prone to overlook very diverse P-Ps due to not-current models and profiles.

Second, this method is poorly scalable and poorly applicable on draft or metagenomic assembled genomes, limiting to assess P-P diversity. Lastly, because it was designed to spot P-Ps broadly without considering the distinct characteristics of specific types, it consequently causes inconsistencies when new members are added to already defined types (e.g., during database updates).

We aimed to improve the detection of P-Ps and their subsequent typing. For this, we developed and present here ‘tyPPing’, a user-friendly method tailored for the distinct P-P types. tyPPing is capable of simultaneously detecting and typing the most prevalent P-P types. It searches for conserved P-P proteins, and uses their frequency and distinct repertoires (compositional sets) to accurately separate P-Ps from other MGEs. Moreover, tyPPing assigns confidence scores (high to low), which facilitates the detection of *bona-fide* P-Ps and atypical, debatable cases. We trained tyPPing on our most recent P-P dataset (18) and compared its performance to our first approach (9) using complete genomes of phages and plasmids, and draft genomes of bacteria. Finally, we compared tyPPing to other classification tools, such as vConTACT v2 (25) and geNomad (19), and we discuss the performance of all approaches, highlighting the benefits and downsides in the strategies of detecting P-Ps.

## Results

### Protein profiles specific for phage-plasmids cover phage and plasmid functions

We aimed to develop a method that is specific and fast in detecting P-Ps. For this, we aimed to use hidden Markov models (HMM) built from protein sequences. We focused on sequences of prevalent, curated P-P types (according to our classification system (9)), and selected types that have at least 10 members and experimentally validated cases (proven to be both phages and plasmids). We excluded small and sparsely-related clusters due to the limited number of available sequences or low-homology to experimentally-proven P-Ps. Our selection included P-Ps of AB_1, P1_1, P1_2, N15, SSU5_pCHM2, pMT1, pCAV, pSLy3, pKpn, and cp32 (key characteristics of these types are listed in Table 1). Notably, in our previous study (9), cp32 P-Ps, reported as φBB-1 phages making 11.2 % of the P-Ps, were not assigned as curated since their lytic genes were unknown. Here, we include them (see Methods, Figure S1), as in a recent study the function of genes involved in the virion formation have now been confirmed and annotated (26).

**Table 1:** Characteristics of the 10 well-defined P-P types. This includes reported examples, along with their virion morphologies, taxa of (frequent) host if P-Ps were found in host of different species, number of conserved protein families (computed by PPanGGOLiN (32)), thresholds for tyPPing and size ranges.

| P-P type | Key example | Virion morphology (of example) | Taxa of host (most frequent) | Conserved protein families (count) | Min Proteins (cutoff) | Composition sizes (cutoff for missed proteins) | Size range (kb) |
| --- | --- | --- | --- | --- | --- | --- | --- |
| AB_1 | SSU5 (distant) | Siphovirus | *Acinetobacter baumannii* | 94 | 55 | 62 – 94 (7) | 101-122 |
| cp32 | φBB-1 | Myovirus (elongated head) | *Borrelia burgdorferi* | 32 | 18 | 24 – 32 (6) | 28-33 |
| N15 | N15 | Siphovirus | Enterobacteriaceae (*Klebsiella pneumoniae*) | 43 | 7 | 16 – 43 (9) | 40-66 |
| P1_1 | P1 | Myovirus | Enterobacteriaceae (*Escherichia coli*) | 71 | 49 | 43 – 71 (11) | 72-125 |
| P1_2 | D6 | Myovirus | Enterobacteriaceae (*E. coli*) | 84 | 46 | 71 – 84 (17) | 79-104 |
| pCAV | SSU5 | Siphovirus | Enterobacteriaceae (*K. pneumoniae*) | 73 | 30 | 42 – 73 (3) | 103-118 |
| pKpn | SSU5 | Siphovirus | Enterobacteriaceae (*K. pneumoniae*) | 89 | 38 | 75 – 89 (2) | 99-125 |
| pMT1 | SSU5 | Siphovirus | Enterobacteriaceae (*Yersinia pestis*) | 92 | 50 | 47 – 92 (6) | 85-115 |
| pSLy3 | SSU5 | Siphovirus | Enterobacteriaceae (*E. coli*) | 95 | 47 | 60 – 95 (3) | 85-134 |
| SSU5_pHCM2 | SSU5 | Siphovirus | Enterobacteriaceae (*Salmonella enterica*) | 90 | 71 | 81 – 90 (3) | 94-126 |

From the 10 P-P groups, we extracted conserved gene families, aligned their protein sequences, and generated HMMs from the multiple sequence alignments (see Methods, Figure 1A). The number of P-P genomes analyzed per P-P type ranged from 14 to 122, resulting in total of 763 HMMs with 32 HMMs for cp32 P-Ps and 95 for pSLy3 P-Ps (Figure 1A, Table 1). We evaluated the profiles’ characteristics, including detection range, diversity and specificity. On average, we found that 69.1% ± 7.7% (mean of coefficient of variance = 16.7%) of genes are conserved per P-P (Figure 1B). To assess profile diversity, we calculated a Neff value pre profile that represents the number of effective sequences in the alignments (27, 28). While Neff values close to 1 indicate highly similar sequences, higher values indicate more diversity. The mean Neff across all P-P types was 1.06 ± 0.06. We found low sequence diversity (Neff = 1.0) for pMT1 sequences that are specific for *Yersinia pestis*. This is likely because *Y. pestis* strains are oversampled and the fact that is a recently emerged species (29). Highest diversity was observed for N15 HMMs (up to Neff=1.4) and cp32 HMMs (all Neff > 1.0) (Figure S2, Table S2).

**Figure 1.**
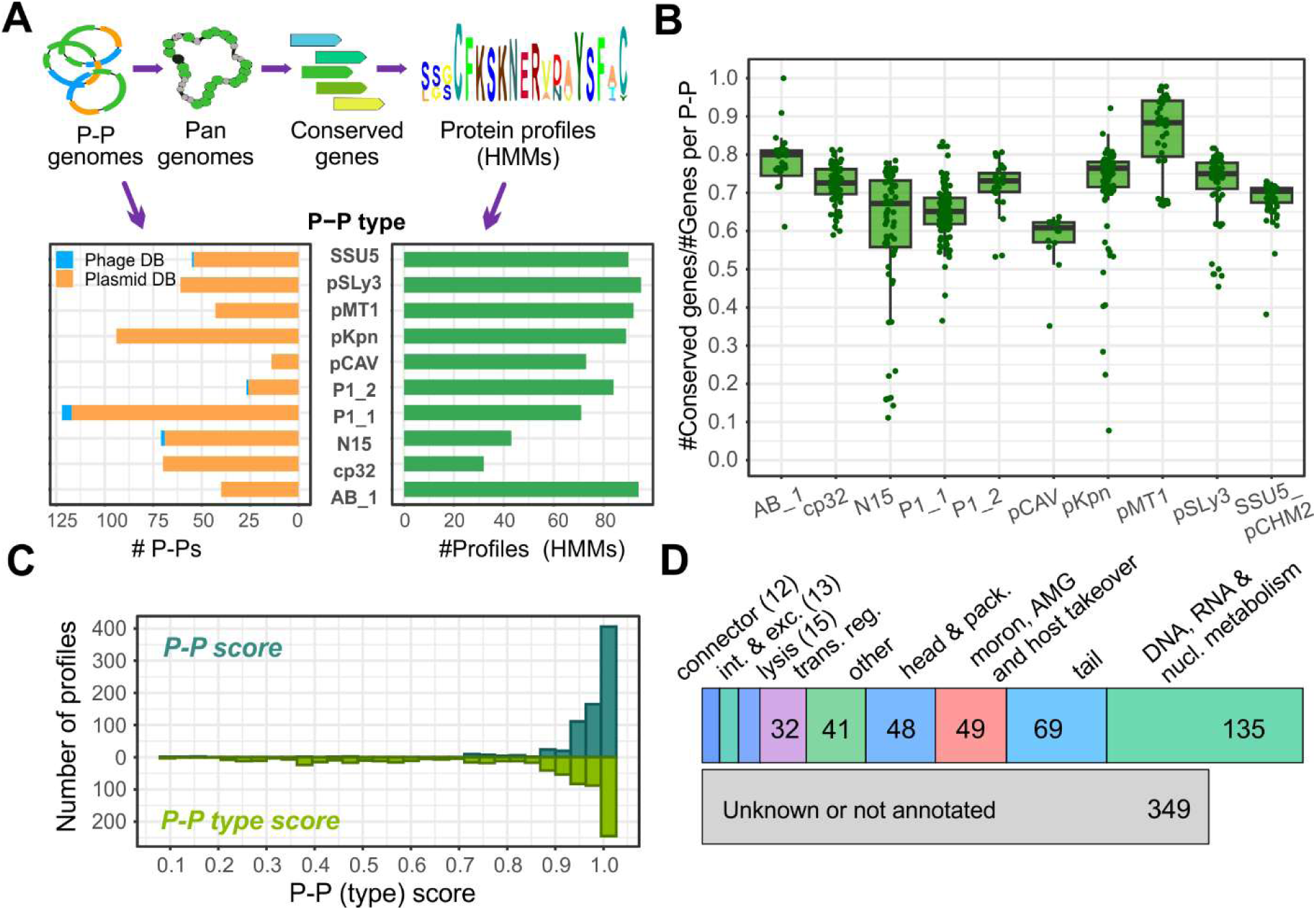
Generation and characterization of P-P protein profiles. (A) P-Ps from 10 curated groups (9) were selected and their pangenomes computed with PPanGGOLiN (32). Conserved sequences (classed as persistent gene families) were extracted, protein sequences were aligned with MAFFT, manually inspected and curated, and used to generate HMMs (see Methods). The number of processed P-P genomes varied per type, and resulted in total 763 profiles (B) Number of conserved genes in relation to the total number of genes per genome (across all P-P types). (C) P-P (dark green) and P-P-type scores (light green) were computed by counting matches of profiles against P-Ps, P-P types, phages and plasmids (see Methods). The two scores range between 0 (unspecific) and 1 (highly specific), and differentiate P-Ps from phages and plasmids or P-P types from other P-Ps. (D) Representative sequences of the 763 P-P profiles were annotated using PHROGs.

Next, we tested the specificity of the HMMs using two scores. The first, named P-P score, was used to differentiate P-Ps from phages and plasmids, and the second, P-P-type score, to determine its specificity to P-P types. The two scores range from 0 (not specific) to 1 (highly specific). Most of the protein profiles (702/763) achieved P-P scores ≥0.9 indicating very high specificity (Figure 1C). However, only 35.4% of the HMMs from pSLy3, pKpn, and SSU5_pHCM2 reached such a high type specificity (P-P-type score ≥0.9), which is a strong contrast to 76% of the other profiles (Figure S3, Table S2). P-Ps from these three types are all closely related to SSU5 and have high genomic similarities (9) causing their profiles to confidently match P-Ps from all three types. Nonetheless, these HMMs can provide valuable information to distinguish these P-Ps from all other P-P types.

To assess functions of the profiles, we annotated representative protein sequences using PHROGs (30), protein profiles specific for replicases and partition systems (31), and the database of geNomad (19). In total, we annotated 64% (488/763) (Table S2), and essential phage (structure, lysis, connector) and plasmid (replication, partition) functions were covered for all P-P types, albeit with varying annotation rates. In particular, while most P1_1 profiles (79%, 56/71) could be annotated, cp32 profiles were the fewest (19%, 6/32) (Figure 1D, Figure S4, Table S2).

In sum, our P-P profiles match on average 70% of P-P proteins, are highly specific to P-Ps (and their types) and cover essential functions of phages and plasmids. From these characteristics, we concluded that these HMMs will function as signature profiles facilitating P-P detection and typing.

### tyPPing uses patterns of conserved P-P proteins to accurately detect and type P-Ps

We used the 763 protein profiles to develop tyPPing that is a workflow for detecting and typing P-Ps. tyPPing requires a profile-to-protein comparison table (produced with the 763 protein profiles), a protein-to-genome and a genome size table (for details, see https://github.com/EpfeiferNutri/Phage-plasmids). tyPPing first computes two parameters (Figure 2A): (1) MinProteins and (2) Composition. For MinProteins, tyPPing counts the number of highly conserved proteins in the target sequences, and keeps only those that pass a threshold (Table 1, Figure S5-S6). In Composition, all proteins that align to least 50% of the HMM sequences are considered. We observed this to be less stringent than the sequence threshold used in MinProteins (Figure S7). Here, tyPPing keeps only target sequences if it detects protein sets similar to those found in P-Ps. To achieve this, tyPPing compares the set of detected proteins (per target sequence) to those found in P-Ps. Because perfect matches (where all proteins in a set are covered) are too restrictive and can miss genetically diverse cases, we allow for a number of missing proteins per set (Table 1, Figure S8). The thresholds for MinProteins (minimum number of conserved proteins) and for Composition (sizes of the protein sets) are distinct for each P-P type (Figure 2A), and we tested various parameters to identify those that enable specific and robust detection (see Methods, Figure S5-S8).

**Figure 2.**
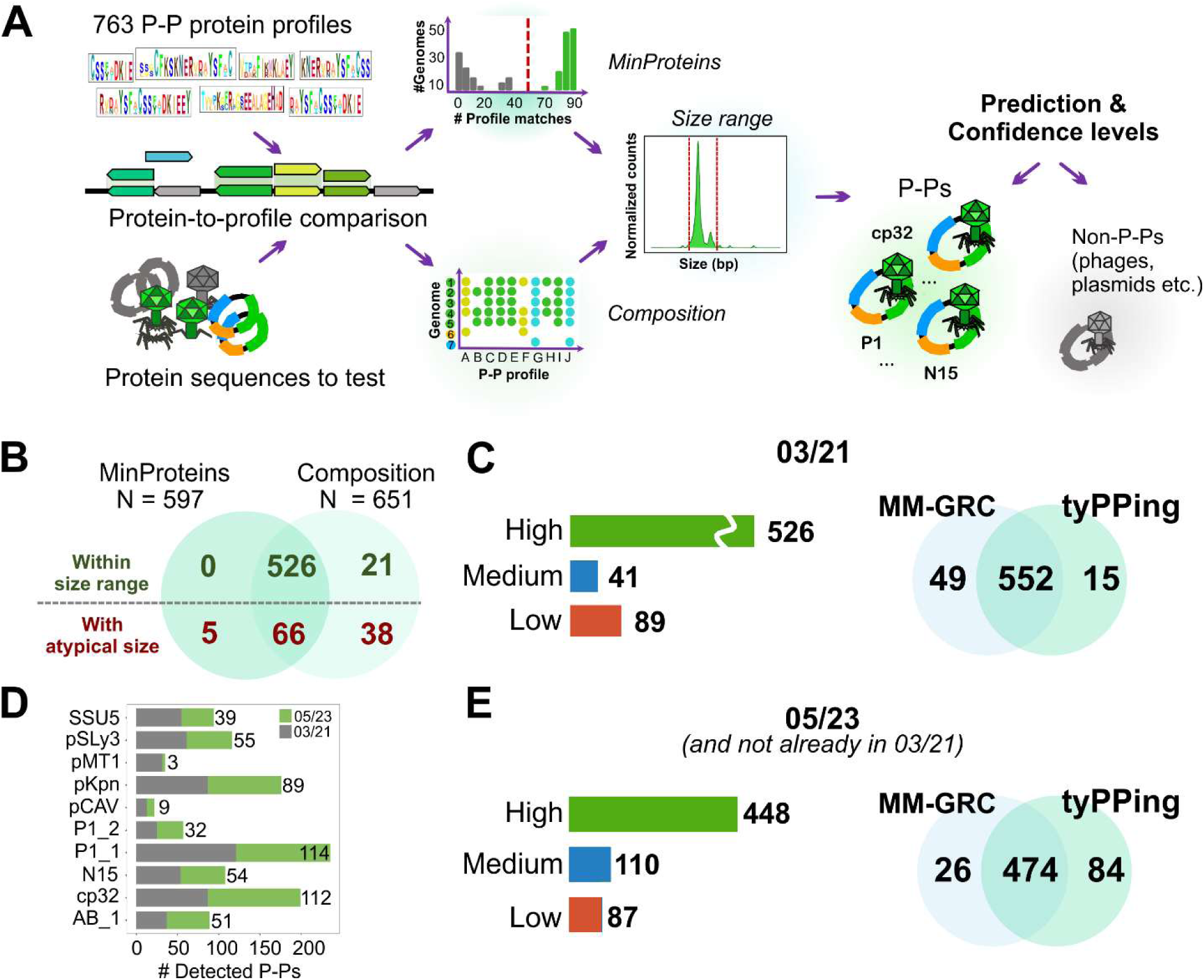
Detection and typing of P-Ps with tyPPing. (A) tyPPing reads in the protein-to-profile table generated by comparing the 763 signature profiles against the to-tested protein sequences, and compare if the sequences meet MinProteins, Composition, and the size ranges of 10 P-P types. To sequences that meet MinProteins or Composition, it assigns confidence levels that depend on the match to the size range (see Methods). (B) tyPPing was employed on the 03/21 dataset, and hits specific for MinProteins and Composition were compared, including the number of elements fitting the type-specific size ranges. (C) Confidence levels of putative P-Ps in 03/21 (bar plot), and intersection of tyPPing and MM-GRC. (D) tyPPing was used to identify P-Ps in 05/23. P-Ps (group-wise) already detected in 03/21 (grey) and in 05/23 (green). (E) Confidence levels of putative P-P sequences of 05/23 (and not already in 03/21) predicted by tyPPing and comparison to MM-GRC predictions.

As a last step, tyPPing assigns confidences to the P-P predictions (high, medium, and low). For this. it uses matches to MinProteins, to Composition and to size ranges of the predicted P-P type (Table 2). We determined the size ranges from curated P-P types, where we excluded elements with atypical sizes (see Methods, Figure S9). A high confidence is assigned to P-Ps that pass MinProteins, Composition and fit the size range of the predicted P-P type. Medium confidence cases are identified by MinProteins and/or Composition, and match the size range with up to 10% variation. A low-confidence case is detected by MinProteins and/or Composition, and has a substantially shorter or longer genome size. We considered sequences with lengths that were either too short (potentially from degradation) or too long (potentially from recombination events or sequencing errors, see Figure S10) to be unlikely P-Ps with functional lytic cycles of phages. This is because they may lack essential genes (too short) or are impaired in packaging the whole genome into the virion (too long). Nonetheless, some of them may represent notable cases of interest (such as defective P-Ps). This is why tyPPing reports these low-confidence cases, and we suggest a closer inspection if they are of particular interest.

**Table 2:** Criteria for confidence levels assigned by tyPPing.

| Confidence | Min-Proteins | Composition | Within size range | Remark |
| --- | --- | --- | --- | --- |
| High | Yes | Yes | Yes | High confidence P-P, meets all criteria |
| Medium | Yes | Yes/No | Yes, or<br>$\pm 10\%$ | Meets MinProteins **and/or** Composition, and fits slightly extended size range |
|  | Yes/No | Yes |  |  |
| Low | Yes | Yes/No | No | As for medium, **but** is substantially smaller or larger |
|  | Yes/No | Yes |  |  |
| Not detected as P-P | No | No | Not tested | Does not meet criteria |

We first tested tyPPing on a database containing over 25,000 complete phages and plasmids (hereafter referred to as ‘03/21’). 656 elements were detected by MinProteins (n=597) and Composition (n=651) with 592 overlapping cases (Figure 2B). 79% were classified with high confidence, 7% as medium, and 14% (n=89) as low (Figure 2B, Table S3). Among the low-confidence elements, seven were identified as too short, while 82 exceeded the expected size ranges. We observed that tyPPing detects sequences that are too-long more often than sequences being too-short. This is because too-short sequences do typically not pass tyPPing’s cutoffs (MinProteins and Composition) and are thus not reported. We also observed that too-long cases often harbor additional sequences with high similarity to plasmids. This suggests that recombination events (between P-P and plasmid) caused these increased sizes (examples shown in Figure S10).

As a next step, we compared tyPPing with our first approach to detect P-Ps, which we will refer as <u>m</u>ulti-<u>m</u>odel <u>g</u>ene repertoire <u>c</u>lustering, or short ‘MM-GRC’ (9). In this method we use phage and plasmid HMMs for functional annotation, random forest models for P-P detection, and employ the gene repertoire relatedness (wGRR) to cluster and type P-Ps (see Methods). The workflow of MM-GRC differs for replicons initially classed as phage or plasmid sequences. Specifically, phage sequences are screened by plasmid specific HMMs for plasmid-related functions. Plasmids are analyzed by phage-specific HMMs and then classed by random forest predictors (9). We first compared tyPPing’s predictions with those of MM-GRC using the 03/21 dataset. 567 putative P-Ps were detected by tyPPing and 601 by MM-GRC (in total, 616 putative P-Ps) (Figure 2C). MM-GRC uniquely detected 49 cases, with tyPPing classifying 43 of those with low confidence (Table S1). The remaining six did not match threshold of MinProteins or Composition. 15 cases were detected only by tyPPing, all with medium confidence, of which 13 were assigned to cp32 (examples in Figure S11) and two to P1_1 (NZ_CP010146, NZ_LR883966). MM-GRC does not use cp32 specific phage profiles, but relies on the clustering to detect cp32 P-Ps. We noted that the threshold for clustering excluded the detection of these elements. The two P1-related cases (NZ_CP010146 and NZ_LR883966) were only detected by Composition. We found that these two cases have most of the conserved P1_1 proteins, but lack proteins important for virion structure and assembly (Figure S12). They also encode origins of transfer (*oriT)* sequences, a hallmark of elements mobilizable by conjugation. We therefore classify them not as P-Ps, but as P1-like plasmids that probably evolved from P1-like P-Ps (18). Finally, we tested tyPPing on a more recent genome database generated in May 2023 (‘05/23’). This database has 77% more plasmid sequences in comparison to 03/21 containing 38,051 elements labelled as plasmids. We focused on plasmids since the vast majority of P-Ps (90.2 % in 03/21, Table S1) were detected in previous plasmid databases. While tyPPing required 8 minutes to process 05/23 (and additional 104 minutes for the protein-to-profile comparison), MM-GRC needed 25h (using the same machines, 25 CPUs). TyPPing identified a total of 558 P-Ps that are not already present in 03/21 with at least medium confidence (increase by 104% in comparison to 03/21). 448 P-Ps were predicted with high, and 110 with medium confidence (Figure 2DE). 87 cases were predicted with low confidence, of which all exceeded the sizes ranges. 84 elements were detected only-by tyPPing and 26 distinctly by MM-GRC (Figure 2F). 83/84 tyPPing-specific cases were assigned to cp32, underlining again its high sensitivity towards cp32 P-Ps. The remaining one was predicted to be a P1_2 P-P (medium). This element encodes conserved P1_2 genes and lacks many essential phage genes (Figure S13) suggesting that it is not a P-P, but a P1-like plasmid. Of the 26 cases distinct for MM-GRC, 21 have atypical sizes (too short) or were classed as low by tyPPing. Three of the other cases matched to protein profiles of AB_1 but did not pass thresholds of MinProteins or Composition. We suggest they are defective P-Ps since they lack many conserved AB_1 proteins (Figure S14). The remaining two, detected-only by MM-GRC, are SSU5-like P-Ps, one of *Buttiauxella* (NZ_CP110081) and one of *E. coli* PI6 (NZ_CP074043) (Figure S15). These sequences did not pass MinProteins and Composition thresholds.

In sum, tyPPing detects efficiently P-Ps of different types and provides confidence scores for the predictions. In comparison to MM-GRC, it reaches >99% sensitivity, >99% precision, and despite a few misclassifications (in total three false-positives in 03/21 and 05/23), it is capable to curate P-P datasets by reporting low-confidence cases.

### P-P detection advances with distinct classification strategies

In previous work, we separated well-related P-Ps, which cluster with experimentally-characterized examples, from sparsely-related groups and distinct individuals (singletons) (9). The well-related types are detected by tyPPing, whereas MM-GRC also identifies the other types (Figure 3A). We wondered if current classification tools, such as geNomad and vConTACT v2, could be used to improve P-P detection specifically, focusing on aspects like robustness, P-P typing and the potential to spot novel P-Ps.

**Figure 3.**
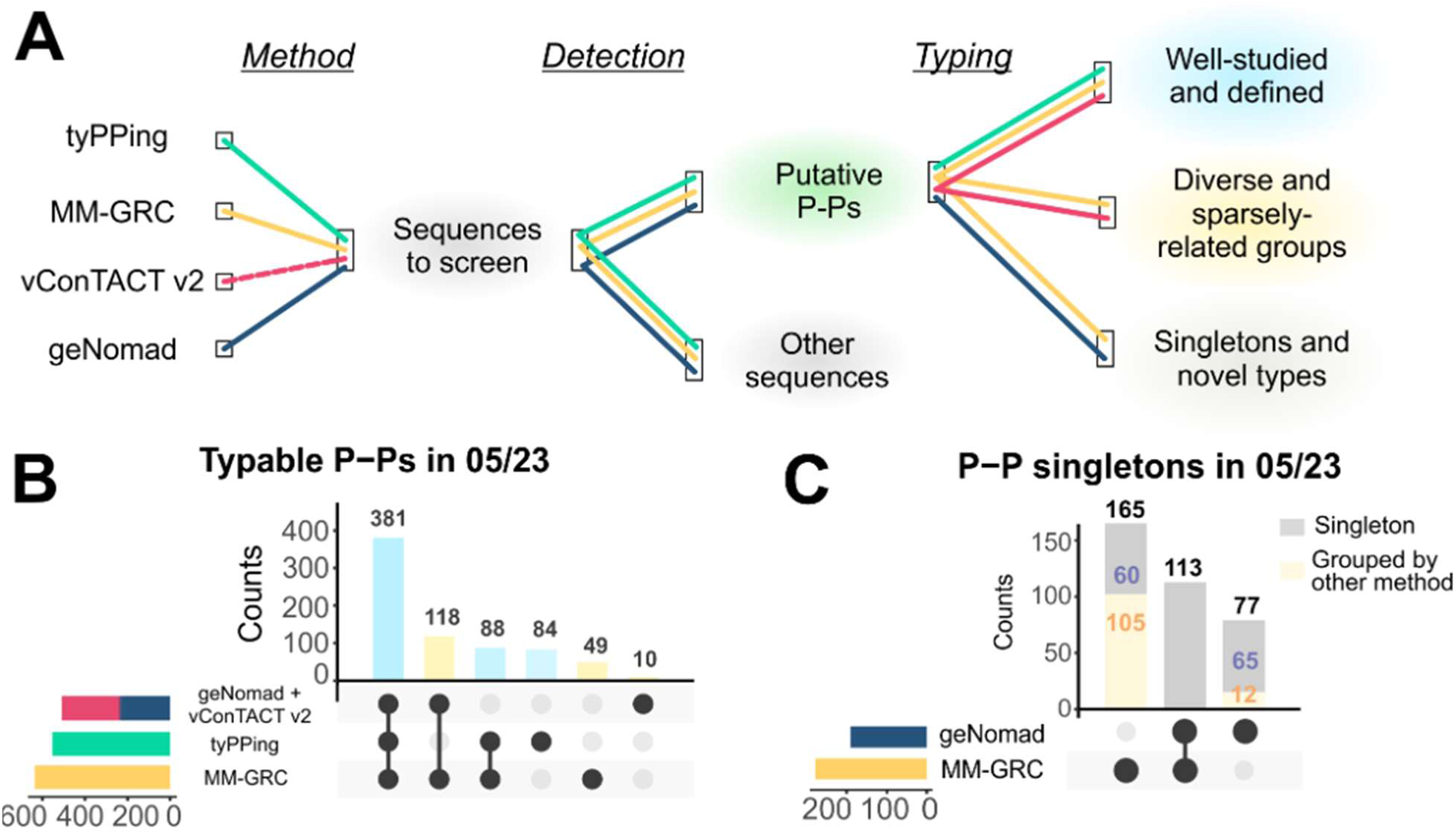
Detection of P-Ps using distinct strategies. (A) P-Ps are categorized in well-defined (blue), sparsely-related groups (yellow) and singletons (grey). Specific methods such as tyPPing, vConTACT v2, geNomad and MM-GRC, were compared in their performances to predict and type P-Ps in 05/23. (B) Counts of P-Ps, well defined types (blue) and sparsely-related groups (yellow), identified by tyPPing, MM-GRC and geNomad combined with vConTACT v2. P-Ps detected by any method as singletons were excluded. (C) Counts of P-P singletons detected by MM-GRC and geNomad (grey bars). Some P-Ps were detected as singletons, but were grouped by the other methods (yellow bar, orange numbers).

We first tested geNomad, which types sequences either as phages, as integrated prophages or as plasmids (19). We analyzed all plasmids of 05/23 and considered elements classed as phage-positives as P-Ps. We excluded those identified as integrated prophages as, per definition, they are not P-Ps. In total, geNomad required 6 hours and 17 min (using 25 CPUs) for the 38,051 sequences and detected 807 putative P-Ps (Table S1). Since geNomad does not group sequences, it cannot type P-Ps. Notably, geNomad predicted 19.5% to 30.8% of the P-Ps detected by MM-GRC or tyPPing, respectively, as plasmids and/or integrated prophages (Figure S16A).

Next, we tested vConTACT v2, a guilt-by-association classifier (25). vConTACT v2 clusters sequences by computing gene-sharing networks with a reference dataset. We used the 1416 P-Ps of 03/21 (18) as the references, and sequences that co-clustered with them were considered as putative P-Ps. In total, vConTACT v2 required 168 hours and 41 minutes for 05/23 (with 25 CPUs), and clustered 1480 cases with the P-Ps (Figure S16B). We inspected these cases and noted that 759 (51.3%) are plasmids that clustered with a few putative P-Ps having similar genes. We tested a stricter clustering analysis, by excluding overlapping and weakly associated cases (see Methods), which, however, did not resolve the plasmid and P-P clusters. This suggests that if solely-used, vConTACT v2 detects a high number of plasmids as P-Ps (making >50% false-positives).

To mitigate limitations of geNomad (inability to type P-Ps) and vConTACT v2 (high number of false-positives), we devised a combined approach. For this, we first used geNomad to separate plasmids and putative P-Ps, and then applied vConTACT v2 for the P-P typing. This approach detected 381 P-Ps of the well-related types (68% of the tyPPing-positives), and assigned them consistently as tyPPing (Figure 3B). 172 P-Ps (identified by tyPPing and MM-GRC) were not detected, and were classed either as outliers (by vConTACT v2) or not as phages (by geNomad). Moreover, geNomad/vConTACT v2 assigned 118 P-Ps to the sparsely-related P-P groups consistently as MM-GRC. While MM-GRC placed additional 49 P-Ps, geNomad/vConTACT v2 clustered 10 cases into distinct groups.

Lastly, we tested the methods’ ability and specificity in the detection of P-Ps that do not cluster with other P-Ps, which we will refer as singletons. These P-Ps may represent novel P-P species or types, making them intriguing for understanding P-P diversity and phylogeny. vConTACT v2 and tyPPing do not detect them, as vConTACT v2 relies on a reference dataset, while tyPPing detects and types simultaneously. Thus, we considered P-Ps that were predicted by MM-GRC and geNomad, and counted them as singletons if they do not group with any other P-P. By this, we detected a total of 355 putative singletons, of which, however, some P-Ps had conflicting classifications. While one method detected them as singletons, another one clustered them with other P-Ps. For instance, MM-GRC identified 165 P-P singletons, but 105 clustered with other P-Ps by the other methods. Conversely, of the 77 P-P singletons specific for geNomad, MM-GRC grouped 12 of those with other P-Ps (Figure 3C).

In sum, tyPPing outperforms any other method in terms of efficiency, speed and sensitivity to detect well-related P-P types. MM-GRC and geNomad/vConTACT v2 are effective to also spot P-P singletons and weakly-connected P-P clusters.

### tyPPing detects complete and active P-Ps of carbapenem-resistant strains

In a final assessment, we wondered if we can use tyPPing to effectively screen unfinished (draft) bacterial genomes for P-Ps. For this, we modified tyPPing and changed its counting system (of conserved P-P proteins) to a cumulative mode. This enabled P-P detection even if the conserved proteins are split across multiple contigs. However, with this setting tyPPing cannot distinguish multiple P-Ps of the same type (e.g. two P1_1 P-Ps) within a single genome, as counts of conserved proteins would be masked. We assume such events are rare, as same-type P-Ps are likely incompatible (as widely reported for plasmids (22)). Importantly, P-Ps of different types (e.g. P1_1 and N15) are still detected. We tested this version on a collection of draft genomes (short-read assemblies) from carbapenem-resistant *Enterobacteriales* species. We focused specifically on a subset of this collection for which we also had corresponding long-read assemblies. Using tyPPing, we detected all P-Ps that we found in previous work (that also encode ARGs) (9) with one medium- and eight high-confidence cases (Table 3) (14). Moreover, we screened three further strains, and detected two P1-like P-Ps in two *Escherichia coli* isolates, 170D8 (P1_1) and 174J8 (P1_2), and one N15-like P-P in *Enterobacter cloacae* 204G7. We observed that tyPPing’s predictions varied with the quality of the assemblies. Specifically, if sequences were just partly or not assembled, then the sum of detected conserved P-P proteins may not meet criteria of MinProteins or Composition, and thus tyPPing may miss these cases (Table 3).

**Table 3:** tyPPing’s P-P predictions for P-Ps detected in short- and long read assemblies, and complete genomes (resulted from hybrid assembly).

| Source | Strain | P-P type | Confidence<br>short read assemblies | Across 'x'<br>contigs | Confidence<br>long reads assemblies | Across 'x'<br>contigs | Confidence<br>complete | Packaged<br>in virion |
| --- | --- | --- | --- | --- | --- | --- | --- | --- |
| Pfeifer<br>et al.,<br>2022<br>(14) | 163A9 | P1_1 | High | 5 | Not<br>detected | 0 | High | Yes |
|  | 166F4 | pSLy3 | High | 2 | High | 1 | Not<br>detected | No |
|  | 169C2 | N15 | High | 2 | High | 2 | High | Yes |
|  | 169C2 | SSU5_<br>pHCM2 | High | 1 | Not<br>detected | 0 | Medium | Yes |
|  | 170E2 | N15 | High | 2 | High | 1 | High | Yes |
|  | 170E2 | SSU5_<br>pHCM2 | High | 1 | Not<br>detected | 0 | High | Yes |
|  | 171A5 | N15 | High | 2 | High | 2 | High | Yes |
|  | 171A5 | SSU5_<br>pHCM2 | High | 1 | Not<br>detected | 0 | High | Yes |
|  | 211G7 | SSU5_<br>pHCM2 | Medium | 3 | Not<br>detected | 0 | Medium | Yes |
| This<br>work | 170D8 | P1_2 | Not<br>detected | 0 | Medium | 1 | High | Yes |
|  | 174J8 | P1_1 | High | 1 | Not<br>detected | 0 | High | Yes |
|  | 204G7 | N15 | High | 2 | High | 2 | High | Yes |

We wondered if the three P-Ps in the three strains 170D8, 174J8 and 204G7 are functional phages and tested if we can induce them by triggering DNA damage (causing a switch into the lytic cycle). For this, we cultivated the strains, added Mitomycin C (MMC), purified phage particles from the supernatants, digested unprotected nucleic acids (mostly from the bacterial host), and extracted DNA from virions (see Methods, Figure 4A). We then sequenced the extracted DNA by short reads and verified the presence of P-P DNA. For this, we first used the short reads to improve the P-P sequences through a hybrid assembly (see Methods), which resulted in complete, closed genomes (Figure S17) and other phage-related sequences (predicted by geNomad). tyPPing assigned a high confidence to these P-Ps (Table 3). Then, we computed the average read coverages of these assemblies, the ones of the host DNA (that was not digested, background signal), and normalized these values (signal of assemblies to background signal). We found that all P-Ps exhibited considerably higher coverages than the host’s level with 36.7-fold for P1_1 (in 174J8), 5.6-fold for P1_2 (in 170D8), and a 93,000-fold for the N15-like P-P (in 204G7) (Figure 4B). In addition, in 174J8, we detected a 43-kb phage (assigned as vc_2 by geNomad) alongside the P1_1 P-P. This phage is related to HK629 (NC_019711, lambda-like), exhibited high coverage (>10,000-fold) and is a residential prophage that was probably activated by MMC.

**Figure 4.**
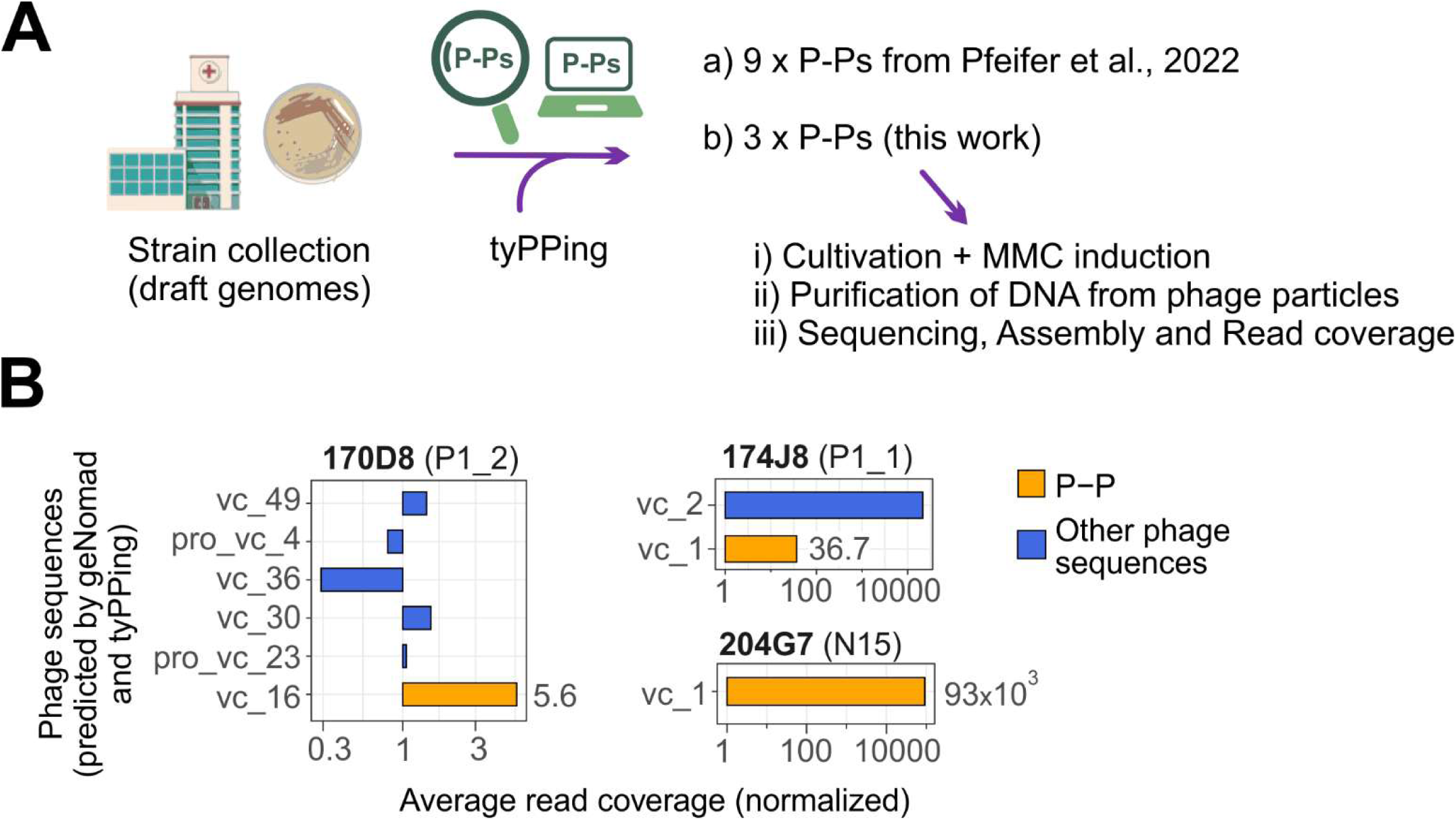
Testing the functionality of P-Ps predicted in draft genomes (by tyPPing). **A.** We modified tyPPing to screen a collection of draft genomes (of carbapenem-resistant species). We detected all 9 P-Ps that we found in previous work (14), and detected three further P-Ps in three other strains (174J8, 170D8 and 204G7). We tested if these P-Ps are responsive to MMC, and packaged into virions upon treatment (see Methods). DNA was extracted from virions, used to conduct a hybrid assembly, and mapped on assemblies to compute the read coverages. **B**. The hybrid assembly resulted in complete P-P genomes (orange), and other phage sequences (blue, predicted by geNomad). The reads from the MMC induction experiment were mapped on all assembled phage sequences. We then quantified the read coverages and normalized these values (of P-P and phage sequences) to the level of the host chromosome.

Taken together, our findings demonstrate that tyPPing effectively detects P-Ps in draft genomes of short- and long-read sequences. While prediction accuracy depends on assembly quality, tyPPing identifies high-confidence P-Ps even when the gnomes are split across multiple contigs. The MMC induction experiments showed that the P-Ps are functional and packaged into virions suggesting them to be active as phages.

## Discussion

P-Ps are widespread in Bacteria, exhibit a high diversity and distinct characteristics (in terms of gene repertoires, hosts, morphologies, sizes) (9). By having genes similar to those of phages and plasmids, accurate P-P detection is a major challenge in microbial genomics. Phages and plasmids recombine, which is promoted by P-Ps (18) and conclusively generates a spectrum of hybrid or chimeric sequences. These events can cause multiple variations of co-integrates, such as fusions between phages and plasmids, or any combination of P-Ps with other P-Ps, plasmids, or phages. While these products retain lifecycles of plasmids (replication and transmission over conjugation termed conduction (35, 36)), they are likely impaired in completing a full lytic phage cycle. This is because recombination can either inactivate essential lytic phage genes or result in genome sizes that prevent complete packaging. For example, while P1 tolerates a size increase up to 10% (of its 93 kb genome (7)), packaging fails at larger sizes (37). Moreover, P-Ps themselves may evolve to plasmids, by becoming defective phages, through gene exchanges and the accumulation of loss-of-function mutations. For instance, we found many plasmids that are related to P1 that lack essential phage genes, but encode sequences that render them as transmissible by conjugation (such as relaxase genes and *oriT*) (18). If these P-P-like plasmids emerged recently, they have many genes similar to those found in P-Ps, phages and plasmids, which makes it difficult to distinguish them from true P-Ps or plasmids. This ultimately complicates accurate P-P detection.

Thus, we aimed to create a refined and calibrated method for detecting P-Ps that also considers their specific types, and developed tyPPing. We tested tyPPing exhaustively on various databases, where it consistently demonstrated effective detection of P-Ps, also in comparison to other methods. For instance, tyPPing detected 13 P-Ps (cp32 in 03/21) that were missed by our previous approach (MM-GRC). MM-GRC was initially designed to detect P-Ps, but it exhibits limitations. It uses (random forest) models that were trained to specifically separate P-Ps from other plasmids by relying on the detection of phage features. Phage functions of cp32 genes were just recently described (26), explaining why these sequences were not detected (by MM-GRC). Even after manually including cp32 P-Ps, the clustering of MM-GRC still excluded the aforementioned 13 cp32 P-Ps due to strict cutoffs (that are necessary to avoid false-positives such as P-P-like plasmids). Moreover, MM-GRC detects putative P-Ps that tyPPing classed with low-confidence (14.8% of well-related P-Ps in 03/21). These sequences, either too-long or too-short, are puzzling, as they appear to be co-integrates, shorter variants, or sequences undergoing degradation. We suggest a manual inspection and careful consideration of these cases since without experimental work one cannot accurately ascertain these are functional P-Ps or defective elements that have become P1-like plasmids. Lastly, MM-GRC needs to be applied separately on phage and plasmid sequences, lacks automation (requires numerous, manual steps) and is constrained by a poor scalability (due to the intense clustering). Nonetheless, it correctly recognized five elements that were misclassified by tyPPing.

We compared our methods, tyPPing and MM-GRC, to geNomad and vConTACT, tools that were not specifically made to detect P-Ps. geNomad detects efficiently MGEs (19), but it does not cluster them. We counted elements of a plasmid database as P-Ps if they were classed as phages by geNomad. vConTACT v2 is limited in the scalability (required >7 days to cluster >38 000 sequences of the plasmid database), and predictions strongly depend on the used references. For instance, we noted if a putative P-P (experimentally not confirmed) encodes too many plasmid-like genes, vConTACT v2 co-clusters these cases with plasmid sequences causing a substantial increase in false-positives (>50% of detected elements). If combined (P-P detection with geNomad, typing with vConTACT v2), the grouping was consistent with those of tyPPing and MM-GRC (for well- and sparsely related P-P types, respectively). However, we noted that geNomad annotated 19-30% of P-Ps as plasmids or integrative prophages, which was most evident for cp32 P-Ps (classed as plasmids). We speculate that geNomad was not trained to detect cp32 P-Ps as phages (as reasoned for MM-GRC).

A current drawback in microbial genomics is the lack of robust P-P databases and classifiers (11) limiting our understanding of the roles and diversity of P-Ps. tyPPing bridges this gap by detecting P-Ps in a fast, scalable, and highly accurate manner. Its detection strategy builds on specific patterns (frequency and matches to compositional sets) of conserved, type-defining proteins. A key strength of tyPPing is its ability to both detect and type P-Ps, regardless of a sequence’s prior classification (as a plasmid or a phage). This is a significant advantage over other methods and it enabled us to adapt tyPPing to screen bacterial draft genomes for P-Ps. Specifically, by this tyPPing avoids difficulties that arise if contigs belonging to the same P-P are assigned to different categories (some parts to phage-like, and others to plasmid-like). We suggest that this version also works with metagenome-assembled genomes (typically described as ‘’MAGs’) when the contigs are reliably binned. Moreover, tyPPing should be compatible with metagenomic assemblies, when the assembled contigs are of good quality and high completeness (as they represent single genetic elements). Quality and completeness scores are assessed by dedicated tools such as CheckV (38). Thus, combining tyPPing and CheckV is a promising approach to mine metagenomic datasets for P-Ps, and, ultimately, create dedicated P-P databases. Furthermore, tyPPing assigns confidence levels to its predictions, which enables the detection of uncertain P-Ps. For instance, it spotted 89 low-confidence cases in our current P-P set (of the 1416 elements (9)). This feature is particularly important to generate high-quality datasets as quality, richness, and robustness of P-P databases are crucial, especially, in the development of future tools that benefit from machine-learning models.

We tested tyPPing on draft genomes and detected three high-confidence P-Ps. We noted that the prediction relies on the quality of the assemblies, and requires that the sequences encoding the P-P proteins are properly assembled and detected by the HMMs. We then (tested and) confirmed that these P-Ps are functional (packaged into phage particles upon MMC treatment). Consistent with our previous work (14), we observed (relative) higher DNA levels of the N15 P-P and of a co-residential lambda-like phage in comparison to the two P1-like P-Ps. This suggests differences in burst sizes that may depend on varying sensitivities to the DNA stress and also on the physiological host conditions (39). Timing to switch to the lytic cycle is crucial because a rapid/early transition increases the chances of producing progeny but risks leaving a healthy host environment. In contrast, a late or slow response gives the host more time to cope with the stress, but may lead to severe damage to the phage’s DNA. From our findings, we cannot conclude whether the P1 P-Ps are late/slow responders or if they are not yet fully adapted to the host, which would also result in lower burst sizes. Nonetheless, we speculate that a host cell would benefit from late responding P1 P-Ps, as the host would not only win time to cope with the stress, but would also benefit longer from P-P-encoded accessory genes. P1 is also widely-reported to have a broader host range (infect many different enteric bacteria beside *E. coli* (40)) than other phages due to its variable tail fiber configurations. This may compensate for a slow or less efficient response, as even a small number of progeny can be sufficient to infect a suitable host that is not targeted by fast-responders with a narrow host spectrum. Future experimental work will be needed to prove or disprove these competition strategies.

Currently, tyPPing is capable to detect 10 different (and prevalent) P-P types, which is its primary limitation. We aim to expand its detection range in future works by adding more P-P specific profiles. tyPPing uses a flexible workflow that readily supports the inclusion of further types (proven in this work by including cp32 P-Ps). New P-P types should ideally group with cases that are experimentally proven to propagate as phages and plasmids. Many potential new P-Ps have been described in diverse hosts like Mycobacteria (41), *Bacillus* (42) (including Betatectiviruses (43)), *Clostridium* (44), and *Vibrio* species (45, 46). An intriguing case is *Carjivirus communis* (the prototypic crAssphage), which is found to be a linear P-P, highly abundant in the human gut, and reported to have a broad host range (47). Since the gut microbiome is a significant reservoir for temperate phages (48, 49) and also rich in P-Ps (50), mining gut metagenomic datasets offers a promising avenue for discovering novel P-P types.

Lastly, we placed P-Ps into three categories: well-defined types, singletons, and poorly-connected clusters. The high number of P-P singletons suggests that we currently assess only a fraction of the P-P diversity. A standardized taxonomy is needed to improve our understanding, as even for the well-defined types (with many experimentally validated examples), a comprehensive classification is missing. For instance, despite being studied for decades and with hundreds available genomes, P1, N15, SSU5 and their closely related P-Ps lack a detailed taxonomy. At most, they are described to the genus/species level or just simply referred as “unclassified *Caudoviricetes*”.

This taxonomic gap is even more pronounced among the poorly-connected groups and singletons. While these cases are intriguing, the low number of available genomes and the lack of experimental confirmation imply that significant work is still necessary to characterize them. Consequently, we suggest that the development of a clear taxonomic classification needs to be prioritized. This would not only avoid the misclassification of P-Ps but also facilitate a better understanding of their prevalence, fate, origin, and their association with bacterial hosts.

## Methods

### Collecting sequences of phages, plasmids and phage-plasmids

We collected two genomic databases on different time points, and the sequences were retrieved from the non-redundant NCBI RefSeq database (51). The first dataset was generated in March 2021, contains 21 550 sequences assigned as plasmids, and the complete genomes of 3 725 phages. We will refer to this data as “03/21”. We detected in this dataset 1416 P-Ps (18) (see Table S1). The second dataset, retrieved in May 2023 (”05/23”) contains 38 051 sequences, all labelled as plasmids. In comparison to 03/21, 17 540 sequences are new in 05/23 (Table S1).

### Similarity of mobile genetic elements assessed by gene repertoire relatedness

To compare MGE pairwise, we computed their weighted gene repertoire relatedness (wGRR), as done previously (9, 18, 52). Specifically, to compute the wGRR, we count the number of best bi-directional hits (BBH) between two MGEs, weight them by protein sequence identities, and normalize the score to the protein number encoded by the smallest genome. To obtain the number of BBHs, we compute pairwise protein alignments using MMseqs2 (v.14.7e284) (53), and keep all hits with an E-value under 10^−4^ with >35% sequence identity covering at least >50% for both protein sequences. The wGRR was then calculated with the following formula:

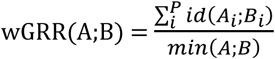

Here A_i_ and B_i_ represent the i-th BBH pair out of a total of P pairs, with the gene count of the smaller element being min(A, B), and the sequence identity between the BBH pair is id(A_i_, B_i_). A wGRR=0 means no matching genes and wGRR=1 means all protein sequences of the smaller element are identical to the BBH in the other element.

### Detection and grouping of phage-plasmids by MM-GRC

To detect P-Ps in 05/23, we used our initial approach to detect P-Ps, which employs <u>m</u>ultiple-<u>m</u>odels approach and combines it with a <u>g</u>ene-repertoire <u>c</u>lustering (MM-GRC) (9). Briefly, we annotated genes with phage-like functions in plasmids using 2583 phage specific HMMs with HMMer v3 (54). A hit was considered as positive, if at least 50% of the profile was covered with a domain i-E-value lower 10^−3^ (as used in MacSyFinder (55). The profiles are grouped in six functional categories: i) structure, ii) lysis, iii) packaging, maturation/assembly and DNA injection, iv) recombination, regulation and DNA metabolism, v) unknown and vi) others. Counts per functional category were used to compute annotation profiles, which served as feature tables for 10 distinctly-trained random forest models. These models were used to predict scores ranging from 0 (plasmid-like) to 1 (phage-like). Scores of the 10 models (one score per each model) were averaged, and elements with values >0.5 were kept. All sequences with sizes >10 kb (smallest reported dsDNA phage) and <300 kb were considered as putative P-Ps. We excluded cases larger 300 kb to minimize false-positives that may arise through co-integration events between mega plasmids and integrative prophages. This step resulted in 887 P-Ps in 05/23 (Table S1). The here-used phage profiles, the random forest models and a customized R script are made available in our GitHub repository (see data availability section).

To type P-Ps we used the wGRR, which we computed between P-Ps from 03/21 and 05/23. Specifically, P-Ps from 05/23 were assigned to types if they had a wGRR of ≥0.5 to a P-P from 03/21 and at least 50% of the genes being homologs. In case of multiple matches, the type of the best hit (highest wGRR) was assigned. Since cp32 P-Ps are not detected by the random forest models as P-Ps (9), we added 39 cp32 P-Ps manually to the 05/23 P-P list using the abovementioned criteria. In total, we detected 926 P-Ps in 05/23 that were not already present in 03/21 (Table S1). Lastly, we placed the 926s P-Ps into well-related (n=498), sparsely-related groups (n=150) and P-P singletons (n=278).

### Curation of the cp32 P-Ps

Circular sequences of sizes of approx. 32 kb were described as P-Ps in *Borrelia burgdorferi* (56). However, MM-GRC did not detect them as such since the protein profiles failed to annotate the phage functions. Thus, in previous work, we placed cp32 sequences (N=87) in a non-curated P-P group, since we could not detect functions typical for phages (9). Nonetheless, in a recent study virion structure proteins (capsid, fibers, tail tube and needles) were described in genomes of five cp32s (NC_012158, NC_012156, NC_012154, NC_012152, NC_012149) (26). We used these cp32 sequences to defined a well-related cp32 P-P group. Briefly, we compared the five cp32 P-Ps with 422 plasmids from *Borrelia* and *Borreliella* (of 03/21) using the wGRR. We kept sequences having at least 50% of genes as BBHs with one of the five cp32s, and a wGRR of 0.6 or higher, resulting in 83 matches (Figure S1). We then considered only elements between 27 kb and 37 kb to exclude too short and too long sequences as they are probably impaired in the lytic phage cycle. In total, 70 elements were kept, and added into a cp32 group in 03/21 (Table S1).

### P-P profiles derived from conserved protein families

We generated 763 signature profiles specific for 10 P-P groups (Table S2). For this, we extracted conserved protein sequences from P-Ps of the well-defined groups (of 03/21): AB_1, P1 subgroup 1 (P1_1), P1 subgroup 2 (P1_2), N15, SSU5_pCHM2, pMT1, pCAV, pSLy3, pKpn, and cp32. Then, in multiple steps, we generated multiple sequence alignments, used these to produce HMM profiles, and then characterized them.

*(i) Extraction of conserved protein families.*

We used PPanGGOLiN (v1.2.63) (32) (default) on sequences of the 10 P-P groups to characterize their pangenomes. PPanGGOLiN utilizes MMseqs2 (53) to cluster protein sequences into protein families with at least 80% identity and a sequence coverage of 80%. Then it calculates a presence/absence matrix of the protein families in genomes, and categorizes them according to their frequencies into persistent (found in most P-Ps), shell (present in an intermediate number of P-Ps), and cloud (found in only a few P-Ps). We selected the highly frequent protein families (persistent or quasi-core) to generate specific P-P profiles.

*(ii) Generation and curation of global multiple sequence alignments.*

We used MAFFT (v7.487) (57) to generate global multiple sequence alignments of each protein family, which we manually inspected, and removed unaligned overhanging regions (solely at the N- and/or C-termini parts).

*(iii) Conversion to HMM profiles and functional annotation.*

Then, we used the curated alignments to generate HMM profiles with HMMER (v3.3.2) (54), and computed a sequence score for each profile, as it is typically done for protein families in PFAMs (58). This score is defined as the minimum score that includes all sequences used in the alignment. We then annotated the P-P profiles using Prokaryotic Virus Remote Homologous Groups (PHROGs) (30), and the database of geNomad (19) by comparing one representative protein sequence to these profiles. Functional categories were assigned for positive hits having E-value ≤ 10⁻⁴ and cover at least 50% of the profile (Table S2, Figure S3).

*(iv) Diversity and specificity of HMM profiles*

To assess profile sequence diversity, we calculated the number of effective sequences, defined as NEFF values (28)]. Briefly, we used HH-suite3 (33) (v3.3.0) to compute these values with the ‘hhmake’ function. NEFF values were extracted from the headers of the resulting ‘*.hhm’* files. NEFF scores measure the average diversity at each position within a sequence alignment. To compute them, the average genetic randomness (Shannon entropy) is computed for small sections of the alignment. For each section a specific Neff value is computed that is the exponential of the Shannon entropy. The final NEFF score is the average of all individual Neff values per alignment.

To assess the specificity of the HMMs, we counted the numbers of P-Ps, phages and plasmids matching them, and computed two specificity scores: (2) P-P score and (3) P-P-type score (Table S2 and Figure S2). A positive hit was assigned if a profile matches a protein sequence, and passes the sequence score. If a protein sequence matched multiple HMMs, we counted only the profile with the best hit (lowest E-value) per protein sequence. The P-P score was computed as follows:

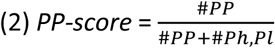

#PP is the number of P-Ps detected (by this profile), and #Ph,Pl represents the total number of phages and plasmids hit (by the same profile). Scores close to 1 suggest high specificity for P-Ps, whereas values closer to 0 indicate a low specificity.

To compute the P-P-type score, we applied the following equation:

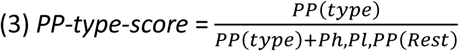

#PP(type) is the number of P-P genomes of a specific type, that were detected by the profile (passed the sequence threshold). #Ph,Pl,PP(Rest) is the number of profile matches against phages, plasmids and other P-P types. A score close to 1 indicates a high specificity in term of identifying the correct P-P type, and close to 0 indicates the opposite.

### Detection of P-Ps and confidence scoring of P-Ps with tyPPing

We used the P-P type-specific HMM profiles to develop tyPPing. Details on how to use it, along with its full documentation, are available in our GitHub repository (https://github.com/EpfeiferNutri/Phage-plasmids). We used sequences of phages, plasmids and P-Ps of 03/21 to define specificity thresholds (Figure S5-S9), and sequences of 05/23 to validate its performance. To use tyPPing, a profile-to-protein comparison table (using all 763 profiles) needs to be computed for protein sequences encoded by a given genome sequence with HMMER (v3.3.2) (54). In addition, information on the genome/contig sizes, and the protein and genome IDs needs to be provided in two tables. Then, by using the provided Rscript, tyPPing predicts if the given sequence is a P-P and assess the confidence level of the prediction. In its workflow, tyPPing first computes two parameters for a given sequence: (1) the number of conserved P-P proteins and (2) which conserved P-P proteins matched this sequence. Next, it compares these parameters to the thresholds of MinProteins and Composition.

Briefly, for MinProteins, tyPPing searches a given sequence for highly conserved P-P proteins and keeps only profile matches that pass the sequence score threshold. If the number of matches is higher than the type-specific cutoff (Figure S5 and S6), the given sequence is kept.

For (2) Composition, tyPPing considers profile matches as positive, if protein sequences cover the matched profiles at least by 50% (Figure S7). Next, tyPPing compares if the detected set of protein sequences matches P-P-specific sets (or compositions). By this, tyPPing retains only sequences having a similar organization as found in genomes of reported P-Ps. To detected also genetically diverse cases, we tested the detection specificity of incomplete sets (up to 75%), i.e. a distinct number of proteins is not detected. We selected those cutoffs that allow the detection of P-Ps and exclude other MGEs (phages, plasmids and P-Ps of other types) (Figure S8). The protein sets, their sizes and allowed number of missing proteins are listed in Table 1 and Table S2.

Lastly, tyPPing assigns confidence levels for the predictions (High, Medium, and Low). For this, it tests a given sequences matches MinProteins, Composition and if it fits a type-specific size range. We determined the size ranges by taking the mean size ± 3x standard deviations of sequences of curated P-P types (excluding P-Ps with atypical sizes, Figure S9). The criteria for the confidence levels are listed in Table 2. Briefly, a high confidence is assigned when a given sequence matches MinProteins, Composition, and fits the size range of the predicted P-P type. A medium confidence indicates that this element is identified at least by MinProteins or Composition, and exceeds the size range up to 10% (of the mean size). A low confidence score is assigned to sequences that substantially deviate from the P-P type specific size range (>10%), and are at least detected by MinProteins and/or Composition.

### P-P detection and typing with geNomad and vConTACT v2

We applied geNomad (v1.7.4, database version 1.5) (19) to all 05/23 sequences labelled as ‘plasmids’ (default parameters). From the virus summary table, we considered 1714 sequences as putative P-Ps since they were identified as ‘phage’ (or ‘virus’) (Table S1). However, it is important to note that geNomad does not perform sequence clustering, meaning it cannot assign types to these putative P-Ps.

To test vConTACT v2 (25), we first clustered all sequences from 05/23 (default settings) with 1416 P-Ps from 03/21 (18). A sequence was considered as a P-P if it grouped with a P-P and was assigned with the same type. We tested two clustering cutoffs. A ‘stricter’ approach, where only cases were considered that clustered clearly with P-Ps (referred as ‘Clustered’ by vConTACT v2). In the second approach, we considered cases that were grouped in overlapping clusters or were categorized as Clustered/Singleton (Clustered, Overlap and Clustered/Singleton are described in the wiki of vConTACT v2. Upon closer inspection, we noted that vConTACT v2 frequently clustered many plasmids with P-Ps even though these plasmids did not encode phage genes, i.e. they are not true P-Ps. The clustering was based solely on the presence of plasmid-typical genes.

We therefore combined geNomad predictions with vConTACT v2’s clustering analysis. Briefly, we used vConTACT v2 to cluster the 1714 geNomad-predicted putative P-Ps with the 1416 P-Ps from 03/21 (after the exclusion of duplicated entries). P-P types were assigned if a sequence grouped with a previously typed P-P (9). A P-P was considered as a singleton if vConTACT v2 could not group it with others.

### Detection of P-Ps in incomplete genome of carbapenem-resistant strains

To screen draft genomes for P-Ps, we modified tyPPing and made its counting of conserved P-P proteins cumulative, i.e. we allowed matches even if they span multiple contigs. Additionally, we excluded counts from contigs larger than 300 kb (as these are likely parts of the bacterial chromosome or mega plasmids), those encoding fewer than 3 protein sequences, and those with less than 10% conserved P-P proteins (relative to all proteins encoded on the contig). The modified version is available in our GitHub repository.

We tested the modified version of tyPPing and compared the results to MM-GRC (9, 14). For this, we screened draft (or incomplete) genomes of the French National Reference center, and detected all P-Ps from our previous work (14), and in three additional strains: a P1_2 in 170D8, a P1_1 in 174J8 and a N15 in 204G7 (Table 3, detailed in Table S3).

We extracted whole genomic DNA of the bacterial strains, and subjected it to sequencing by long reads. Briefly, cells were cultivated in 4 mL LB-Miller medium at 37°C, for ∼16 h (overnight) with shaking at 250 rpm. After a centrifugation step, DNA from cell pellets were isolated with a modified version of the guanidium thiocyanate method (prior to DNA precipitation, the samples were treated with RNase A at 37°C for 30 min) (59). DNA library preparation (SMRTBell Library 10 kb insert size) and sequencing were done at the Biomics sequencing platform of the Institut Pasteur (C2RT) (Paris, France) with the technology of Pacific Biosciences (PacBio). The obtained reads were assembled by flye (v.2.7.1-b1590) (60) with default parameters.

### Mitomycin C induction experiments and purification of phage particles

We tested if P-Ps, that we predicted in the strains 170D8, 174J8, 204G7 are inducible by Mitomycin C. We first cultivated them in 4 mL LB-Miller medium at 37°C for ∼16 h (overnight). Stationary cultures were diluted 1:100 and cultivated in new 4 mL LB-Miller medium at 37°C medium for 1 h. Mitomycin C (MMC) (Sigma-Aldrich, St. Louis, United States) was added in final concentrations of 1 μg/mL. 4 h after (MMC addition), samples of 2 mL were taken for PEG precipitation and pelleted. The supernatant was filtered using 0.22 μm filter and was referred as phage lysates.

### Extraction of DNA from phage particles and sequencing

To purify phage particles, we treated phage lysates with poly-ethylen-glycol (PEG). For this, we added PEG-8000 in a final concentration of 10% (w/v), and NaCl of 0.5 M. The mix was inverted several times, and chilled on ice overnight. Next, virions were pelleted at 60 min and 5,000 × g, 4 °C, and the supernatant was carefully discarded. Pellets were resolved in 100 ul buffer solution (10 mM Tris HCl, pH 7, 10 mM MgCl_2_). To remove bacterial host DNA and RNA, we treated the samples with 2 U Turbo DNAse for 60 min at 37 °C and 5 U of RNAse H. DNase and RNase were subsequently inactivated by adding Proteinase K in a final concentration of 0.1 μg/mL and 0.5% SDS at 62 °C for 20 min. Viral DNA was extracted using a phenol-chloroform method (61). Briefly, an equal volume of phenol:chloroform:isoamyl alcohol (25:24:1, v/v/v) was added to the sample, mixed by inversion, and centrifuged at 10,000 × g for 30 min. The upper aqueous phase was then transferred to a new tube and extracted with an equal volume of chloroform:isoamyl alcohol (24:1, v/v) using the same procedure.

DNA was precipitated by adding 0.1 volumes of 3 M sodium acetate (pH 5.2) and 2.5 volumes of ice-cold absolute ethanol, followed by overnight incubation at −20 °C. The precipitated DNA was collected by centrifugation at 12,000 × g for 60 min at 4 °C, washed with 500 μL of 70% ethanol (centrifuged at 12,000 × g for 15 min at 4 °C), and finally resuspended in Tris-HCl buffer (pH 8.0). DNA concentration was determined using a Qubit fluorometer. For P-Ps from 170D8 and 174J8, Library preparation, sequencing (paired end, 150 bp), and quality checks were performed by Eurofins Genomics (Ebersberg, Germany). For 204G7, these steps were prepared on Biomics sequencing platform of the Institut Pasteur (C2RT) (Paris, France) by short reads (paired-end, 250 bp length). Read sets were made available in the European Nucleotide Archive (see data availability section).

### Hybrid assembly, read processing and mapping

We removed adapters and low-quality reads with fastp (v0.23.4) (default parameters) (62). We then used them together with long-reads (from the PacBio sequencing), and assemble the P-P genomes with Unicycler (v0.5.0) (63). We analyzed the sequences with geNomad (v1.7.4) with default parameters (19), and tyPPing. We then aligned the reads to the assemblies with bwa-mem2 (v2.2.1) (64), and filtered out low-quality alignments with msamtools (v1.1.3) (https://github.com/arumugamlab/msamtools) using the parameters ‘-l 80 -p 95 -z 80 –besthit’. Mapped reads were counted using the msamtools’ profile function with ‘--multi=proportional’. To compute an accurate signal from the host sequences, we first mapped the reads on phage and P-P sequences and collected all unmapped reads (=non-phage) with samtools v1.9 (65). For 174J8 and 204G7, we then aligned these non-phage reads on the long-read assemblies to calculate the average read coverage. In total, 98.4% for 174J8 and 99.4% for 204G7 of the non-phage reads mapped on the long-read assemblies. For 170D8, we achieved a higher read coverage (94.9% of the reads) when we mapped the non-phage reads on the hybrid assemblies, instead of the long-read assemblies, as these were of too low quality.

## Data and Code availability

Rscripts, data tables, sequence, and meta-information to reproduce figures and analysis are available on the GitHub (https://github.com/EpfeiferNutri/Phage-plasmids) and Zenodo (10.5281/zenodo.16616313) repositories.

The raw sequencing reads (short reads from the MMC induction experiments, and long reads from the whole genome sequencing) generated in this study have been deposited in the European Nucleotide Archive (ENA) under the project accession number PRJEB92425, and are also available in the Zenodo repository (for direct access) while being under revision.

Any additional information required to reanalyze the data reported in this paper is available from the corresponding author upon request.

## Acknowledgements

We gratefully thank Marie-Agnes PETIT, and her team at MICALIS, for fruitful discussions, advices and support.

This work was supported in part by the French junior professor chair ANR grant for EP (ANR-22-CPJ1-0041-01) and the Laboratoire d’Excellence IBEID Integrative Biology of Emerging Infectious Diseases [ANR-10-LABX-62-IBEID] and the INCEPTION project [PIA/ANR-16-CONV-0005] for EPCR.

We are grateful to the computational and storage services (TARS cluster) provided by the IT department at Institut Pasteur, Paris, and the INRAE MIGALE bioinformatics facility).

Biomics Platform, C2RT, Institut Pasteur, Paris, France, supported by France Génomique (ANR-10-INBS-09) and IBISA.

## Author contributions

Conceptualization: EP and EPCR

Computational analyses: EP and KI

Biological analysis of the samples: KI and EP

Contributed biological material: EPCR, RB

Sequencing of the samples: EP, KI

Writing—original draft and designed figures: KI and EP

Writing—review & editing: KI, EP, EPCR

All authors revised and contributed to the current draft of the manuscript.

Supervision and funding acquisition: EP and EPCR

---

**Figure S1.**
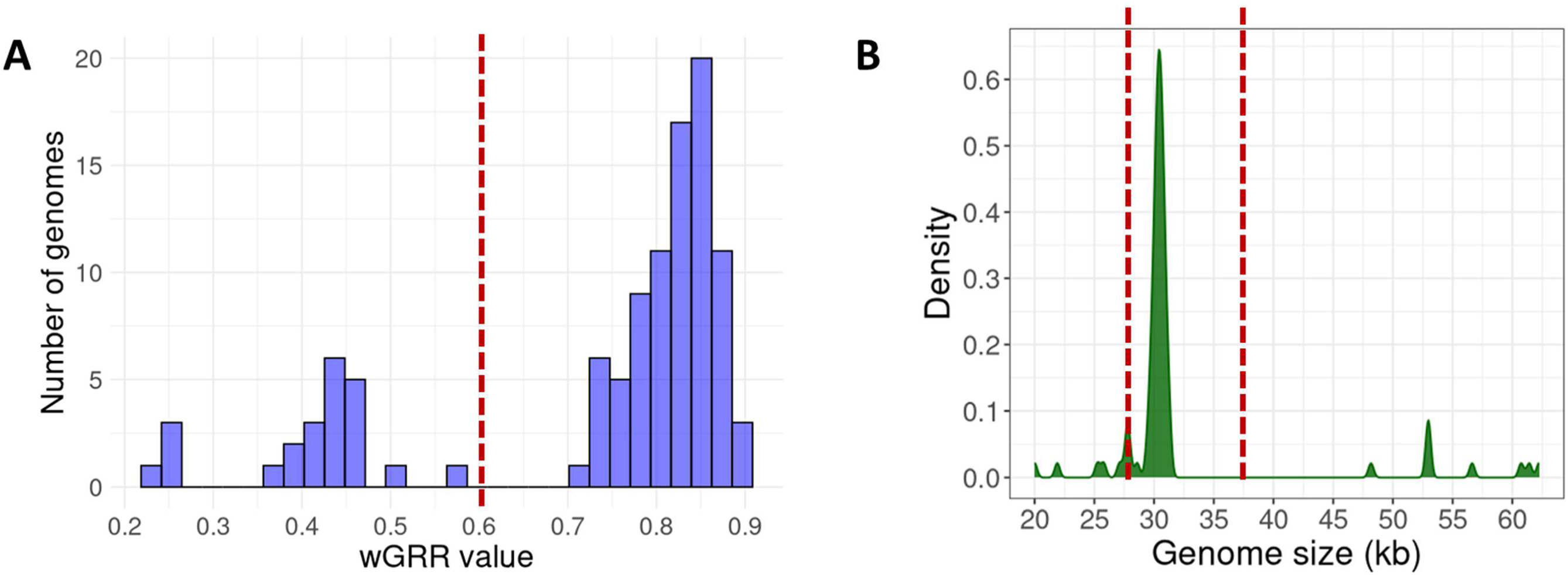
Defining cp32 sequences as a well-related P-P type. (A) Sequences of *Borrelia/Borreliella,* taken from of 03/21, were considered as putative P-Ps if they had homologs (at least 50%) in one of the five cp32 P-Ps (26), and a wGRR >0.6. (B) Some of those elements were substantially larger or smaller than the first reported cp32 P-P (with 32 kb). To exclude co-integrates and fragmented variants, we defined elements with sizes between 27kb and 37kb (red dashed lines) as cp32 P-Ps.

**Figure S2.**
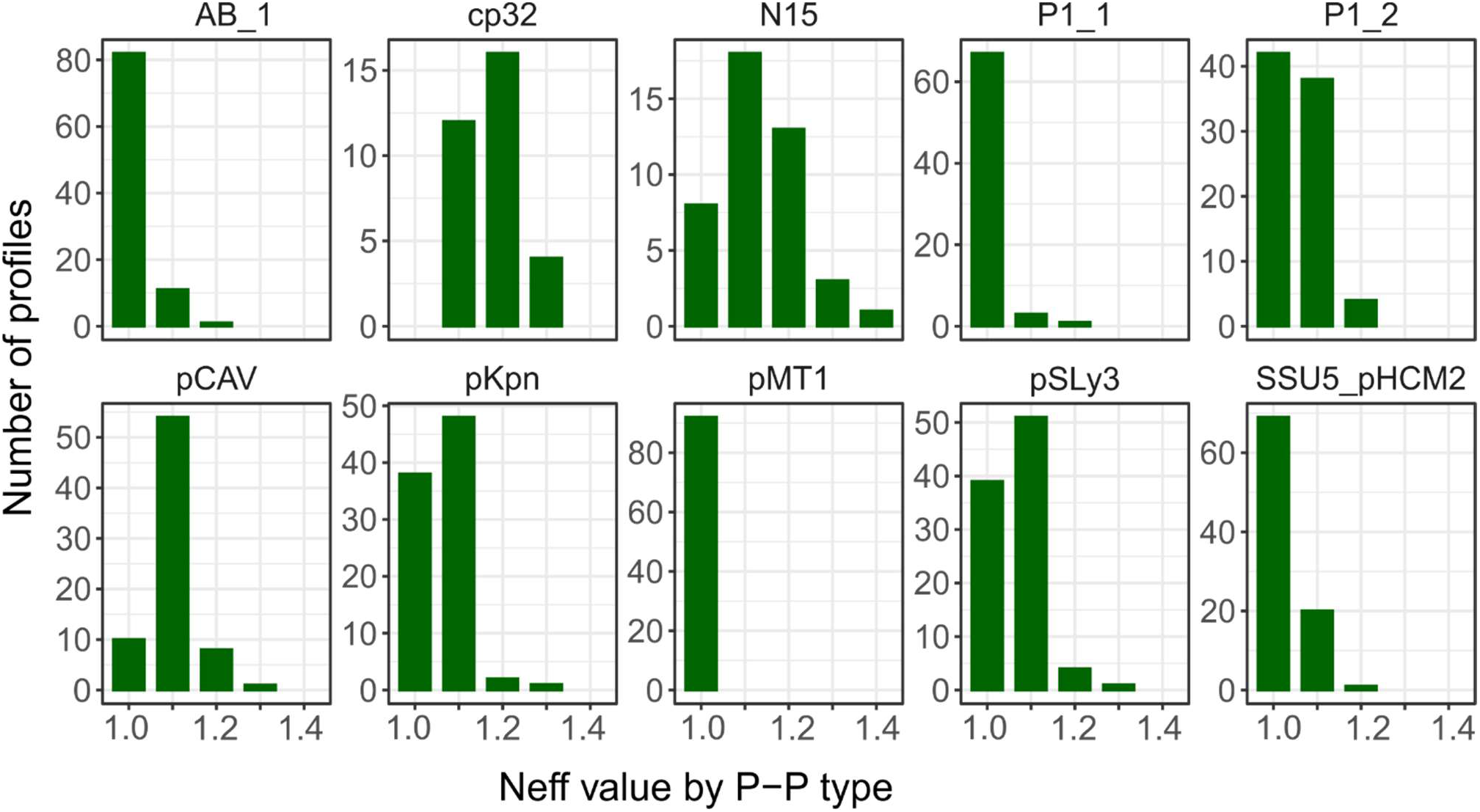
Neff values of P-P profiles across all well-related types. Neff values were computed using HH-suite (33), and indicate the effective sequence diversity of the multiple sequence alignment of the used protein sequences. Whereas Neff = 1 shows low diversity, higher values indicate higher diversity.

**Figure S3.**
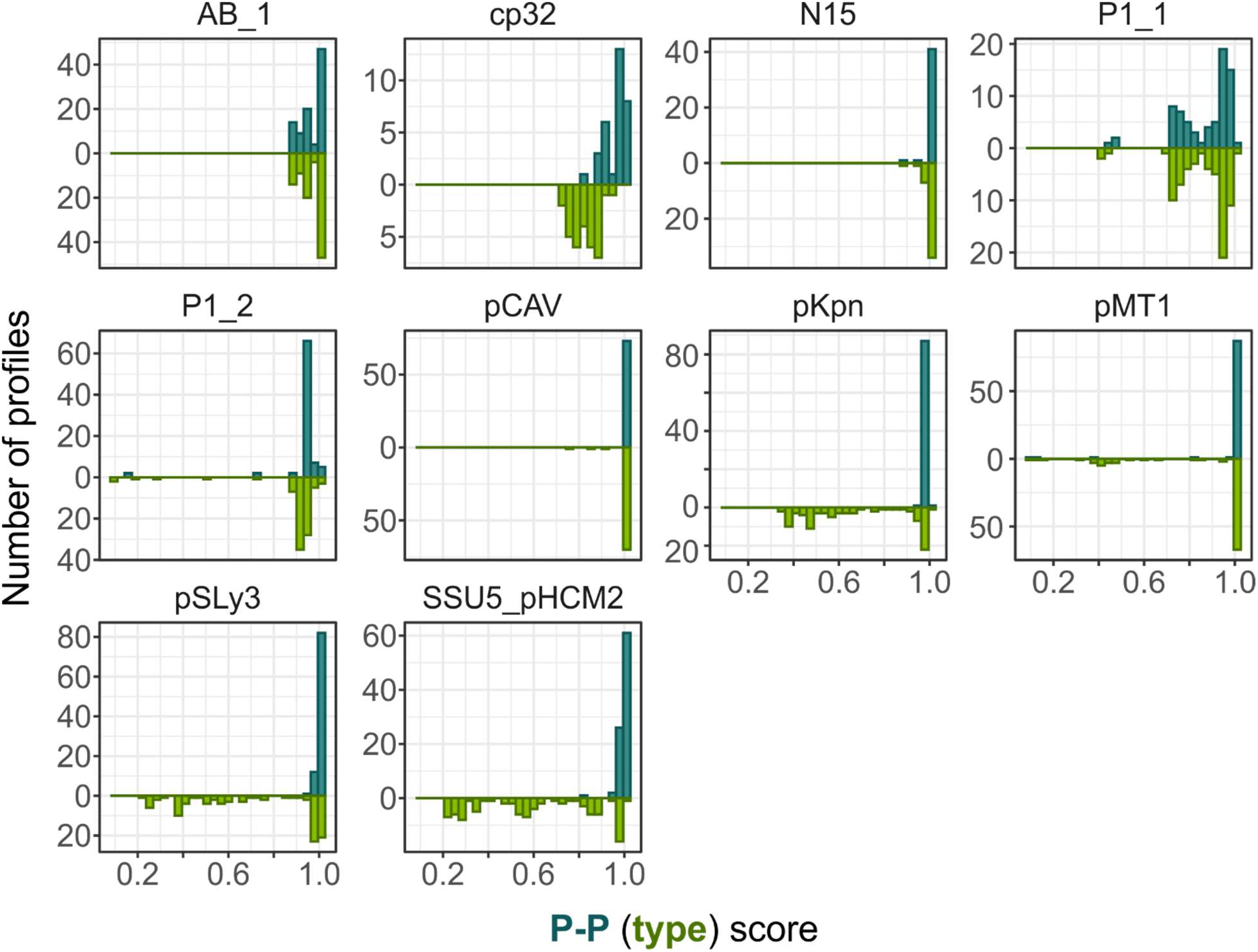
P-P and P-P-type score across P-P types. As in Figure 1C, but group wise.

**Figure S4.**
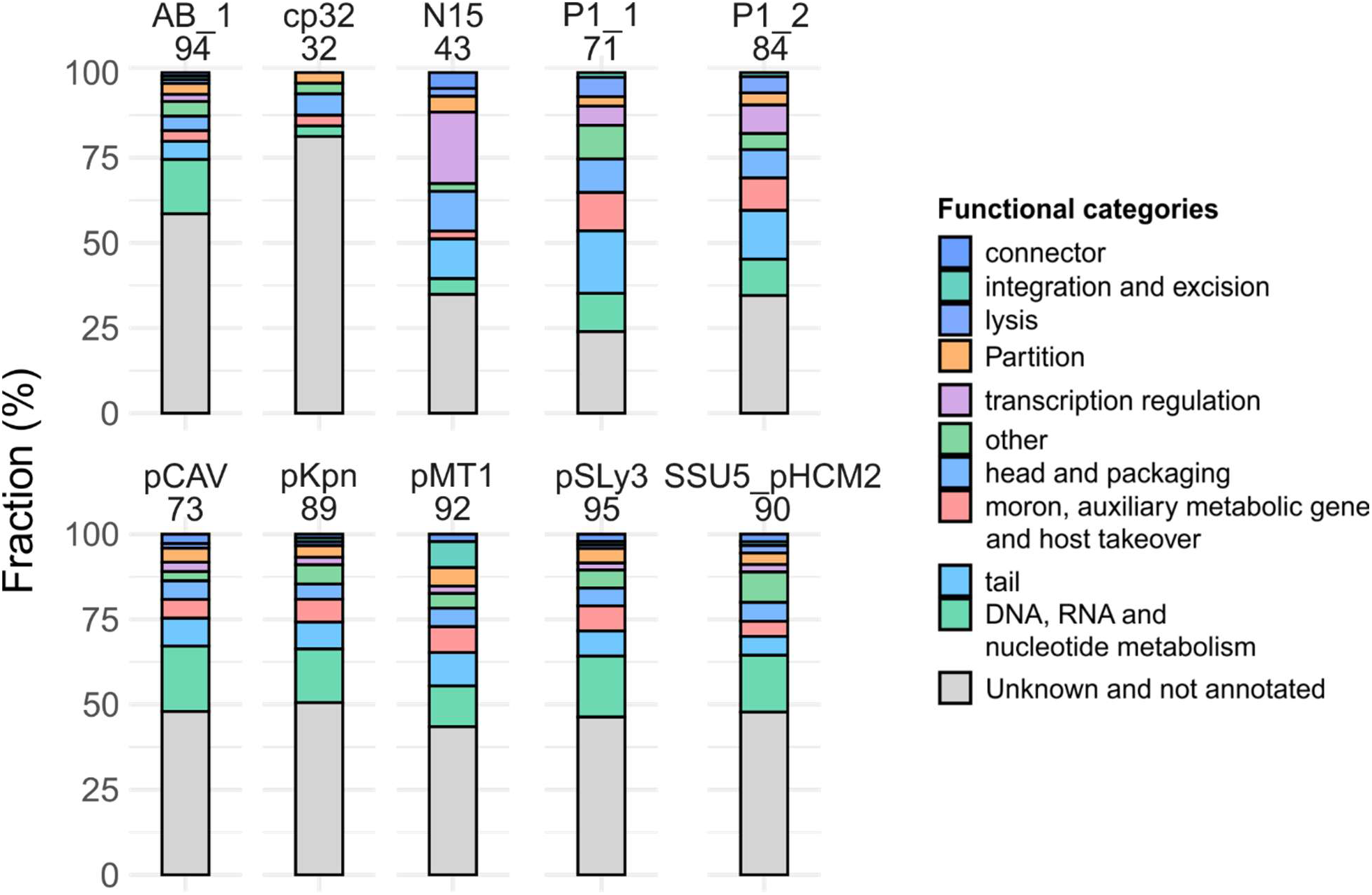
Annotations of the 763 P-P profiles. (as in Figure 1D, but group wise).

**Figure S5.**
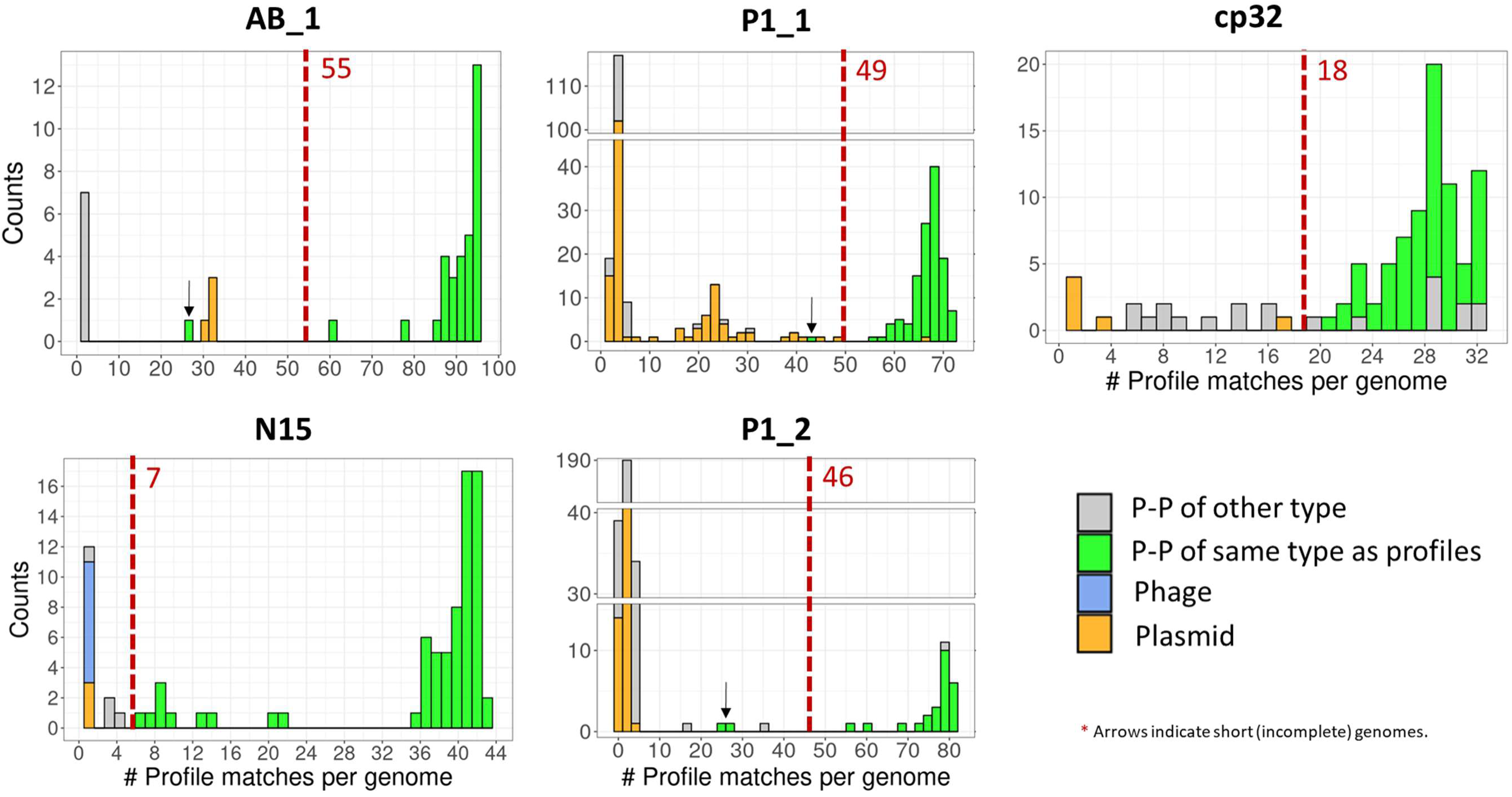
MinProteins thresholds. We used the 763 P-P profiles to search for minimum number of conserved proteins (for AB_1, P1_1, cp32, N15 and P1_2) that allow the differentiation between plasmid, phage and P-P (type). We counted the number of detected genomes (y-axis) and correlated it with the number of profiles that matched these genomes. Chosen cutoffs are shown in red dashed lines.

**Figure S6.**
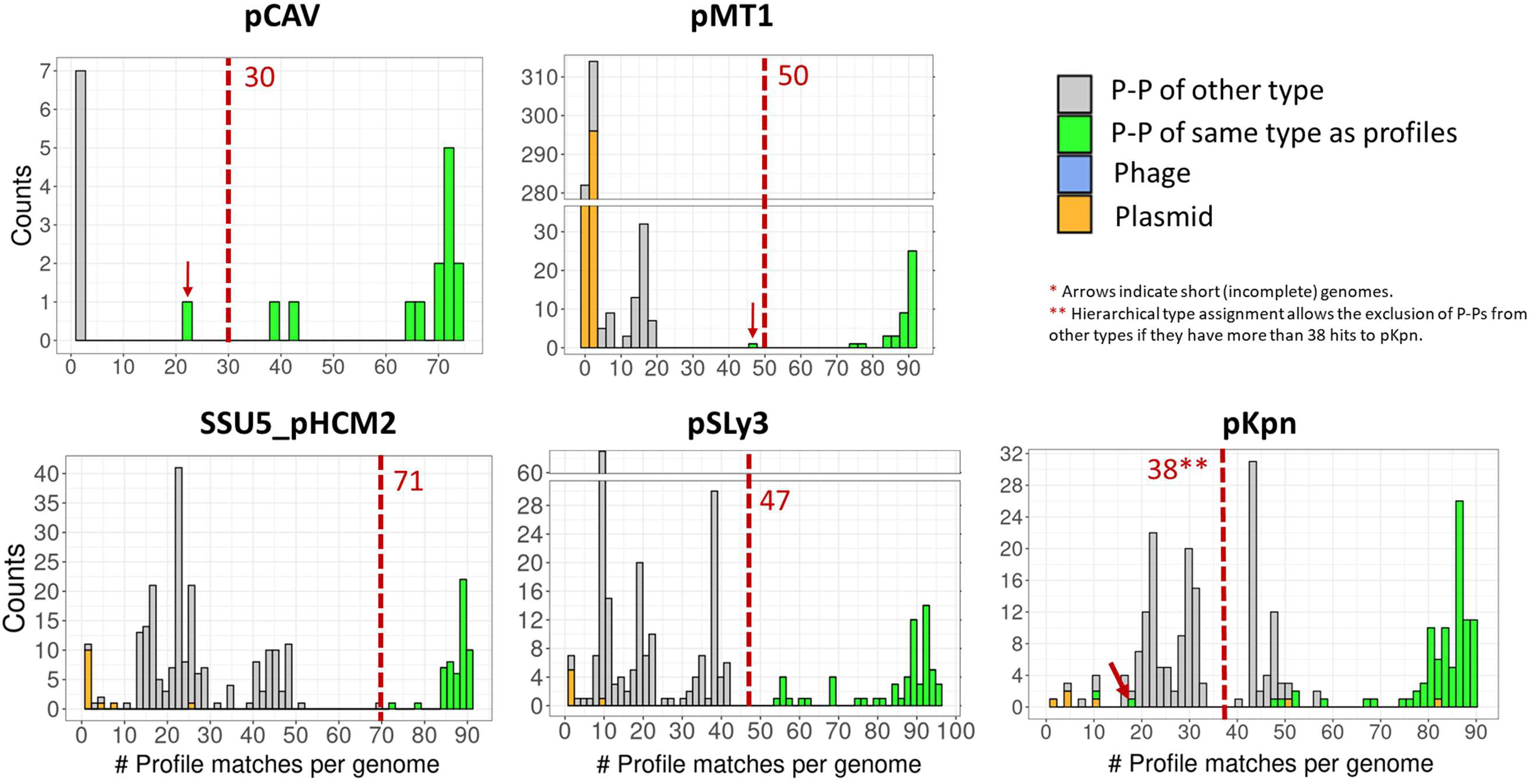
MinProteins thresholds. As Figure S5 but for SSU5-related P-P types.

**Figure S7.**
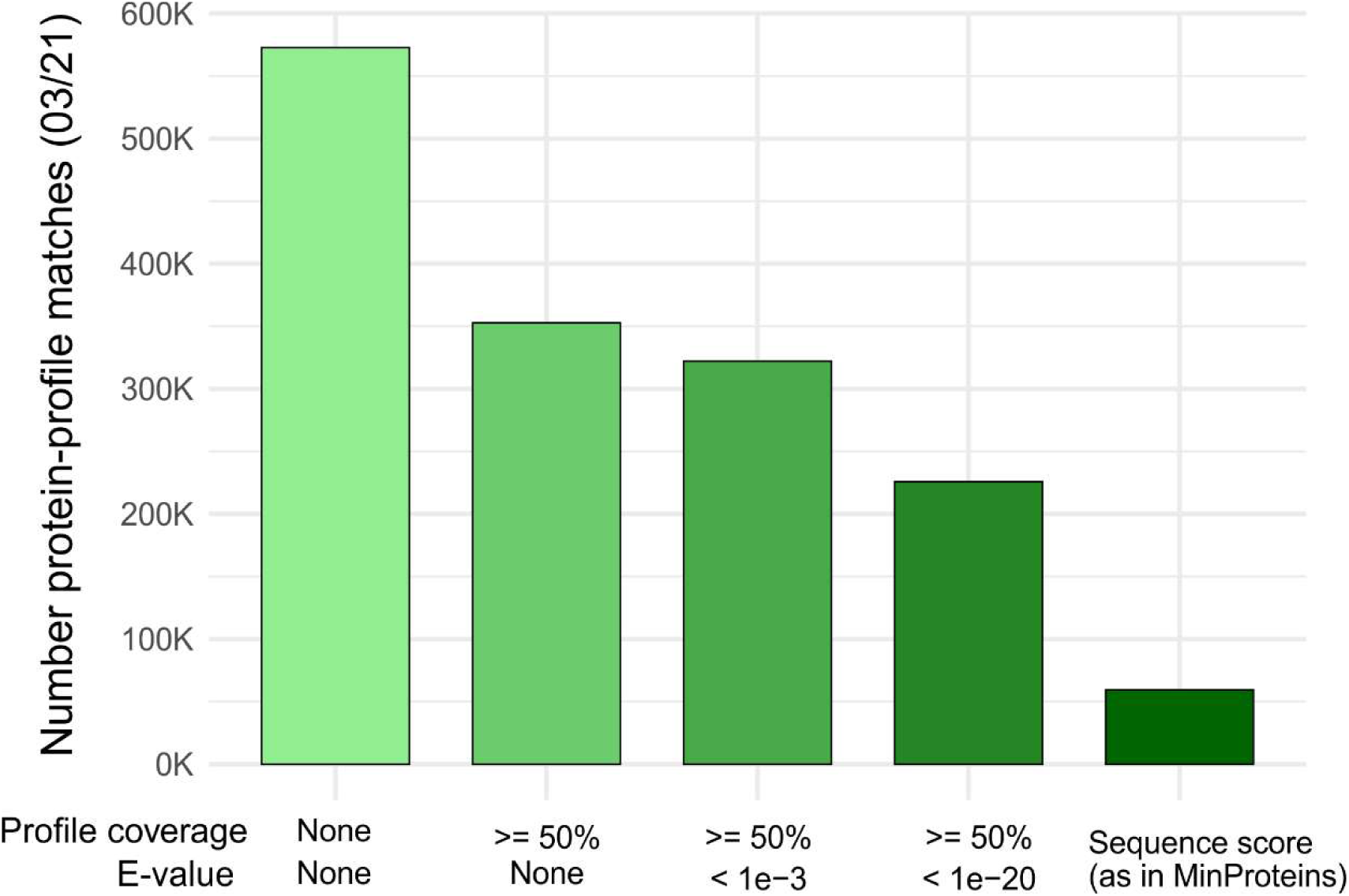
Thresholds for profile-to-protein hits. We evaluated several parameters, such as profile coverage and E-value, to count the number of detected proteins using the 03/21 dataset.

**Figure S8.**
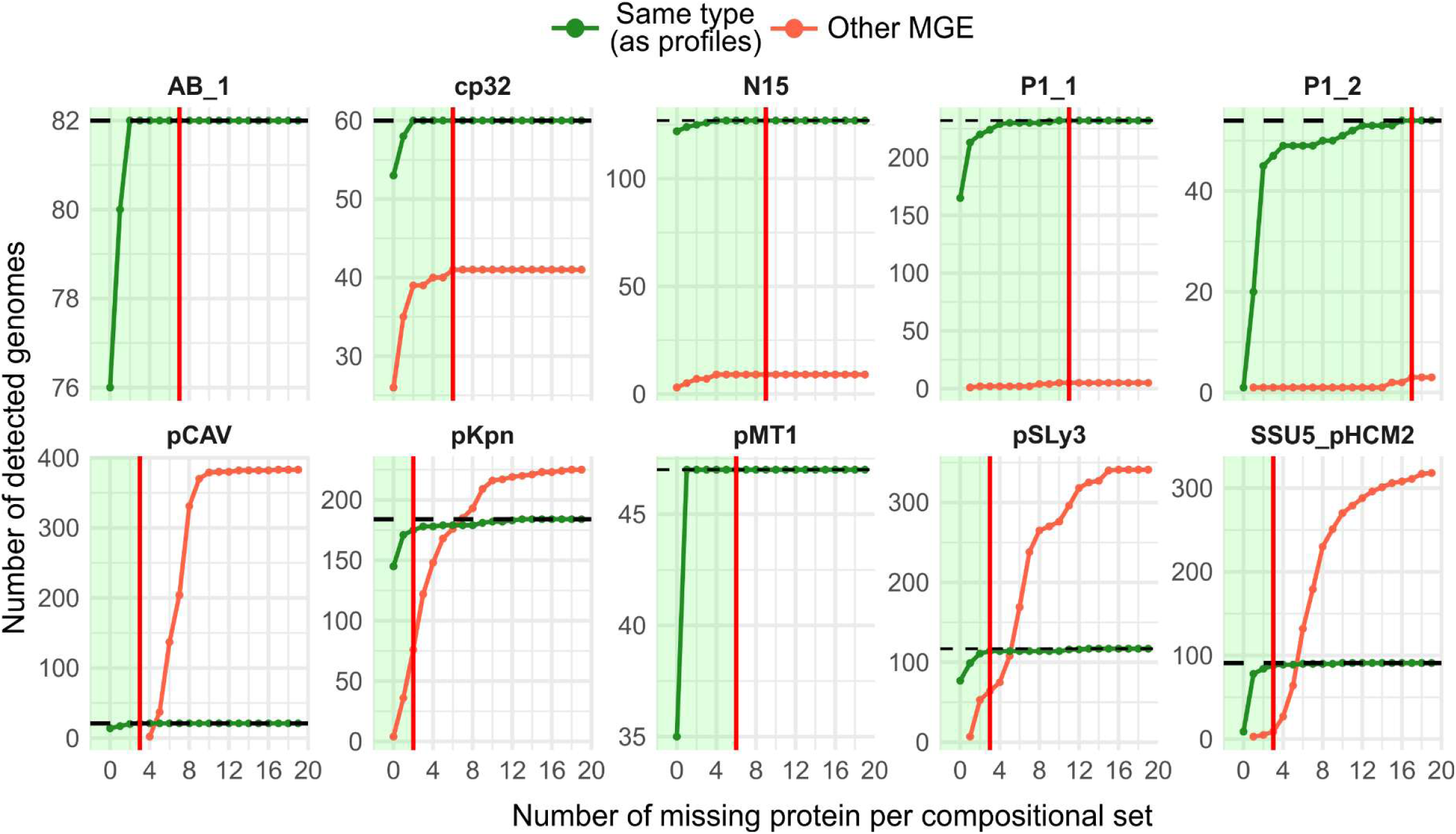
Thresholds for Composition. To count profile-to-protein matches, we fixed 50% coverage (Figure S7). We then varied the minimum sizes of the unique compositions by allowing distinct number of proteins not to be detected (x-axis), and tested the detection specificity, i.e. if a P-P of the same type (green curve), a different type, or a plasmid is recovered (orange curve). Red lines indicate the chosen threshold of allowed missing proteins per P-P type. Notably, to keep robust predictions, and since a high number of missing proteins has a stronger impact on compositional sets with small sizes, the maximum incompleteness was set to 75%.

**Figure S9.**
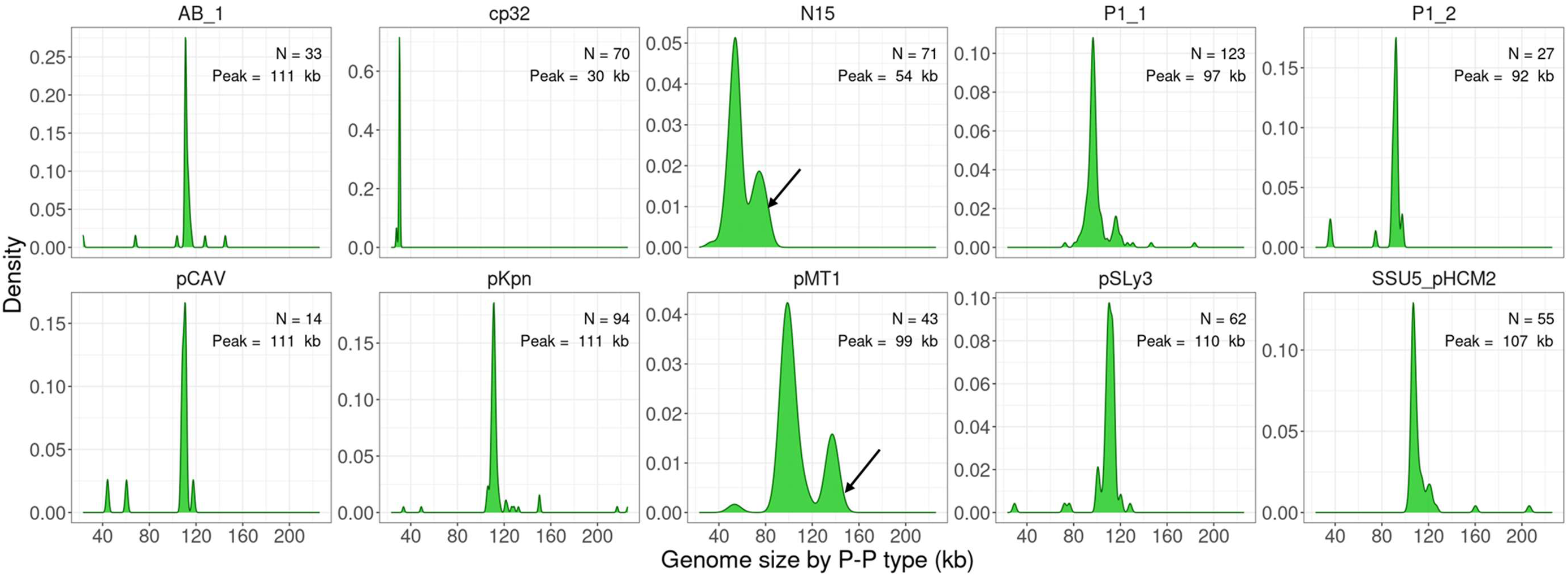
Size distribution of P-P genomes of different types. We first curated the P-P types by excluding sequences with atypical sizes. Specifically, genomes falling outside the major peak and sequences within a shoulder peak (indicated by the black arrows) were not considered in computing the size range. We used the mean size of the curated P-P types, ± 3x standard deviations as the ranges. We found that the unusually long N15 genomes contained long inverted repeats (see Figure S10C for examples) that are reported to be sequencing artefacts caused by wraparound reads (from sequencing linear genomes with the PacBio technology) (34). We found the second population within pMT1 P-Ps (black arrow) to consist of pMT1 P-Ps fused to a plasmid with conjugative genes.

**Figure S10.**
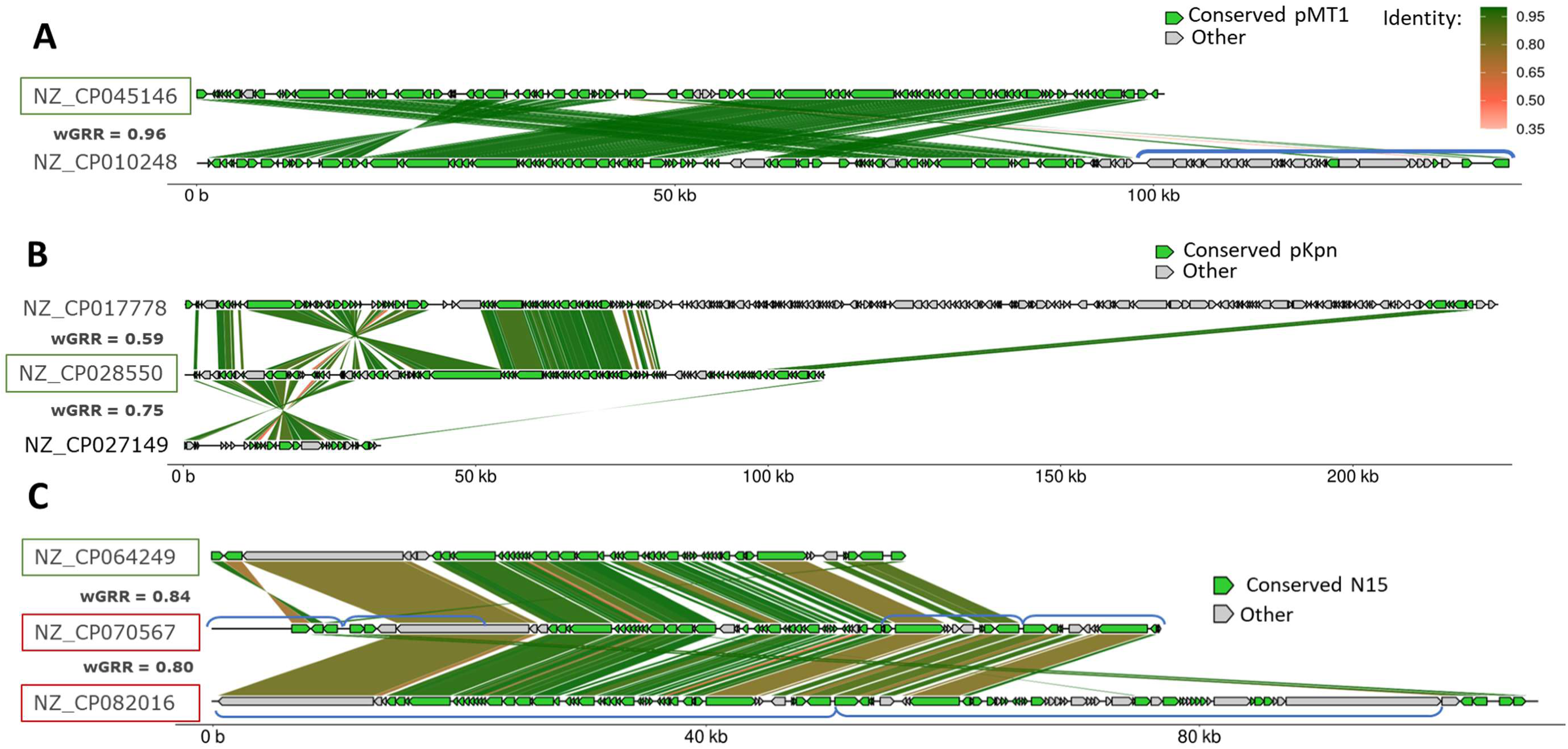
Examples of elements with atypical sizes. (A) pMT1-like P-P (high confident, green box) and a too-long element with additional sequences (indicated with blue brackets). (B) pKpn-like P-P (green box, NZ_CP028550), NZ_CP017778 (first row) is a too-long example and NZ_CP027149 (third row) is a too short example (detected only by MM-GRC). (C) Many of the N15-like cases (n=17, >72 kb) contain long inverted repeats of several kilobases. These repeats likely arose through wraparound reads that are commonly reported for linear elements if sequenced by PacBio (34).

**Figure S11.**
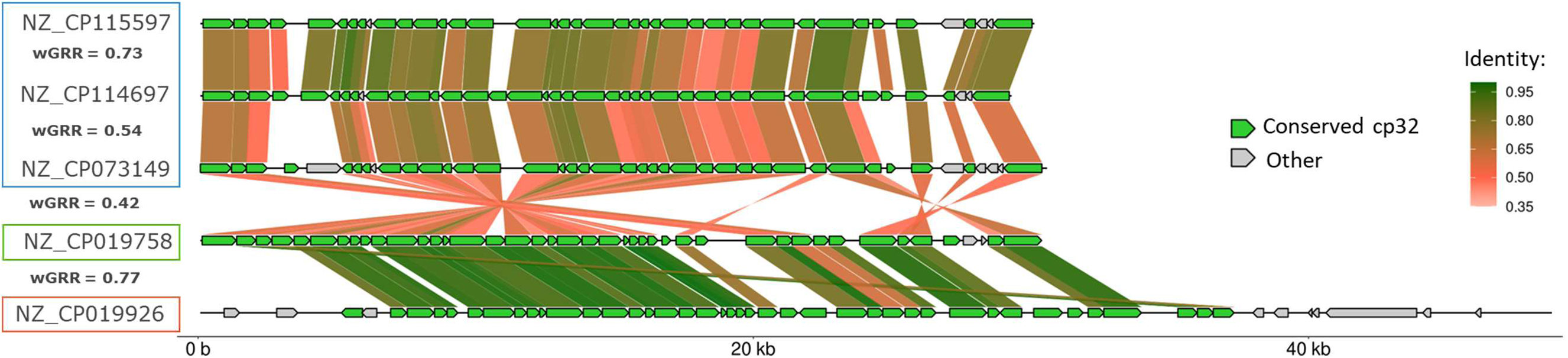
cp32 elements detected as P-Ps or excluded by tyPPing. cp32 P-P with high confidence (green box), an element with a too-long size (low confidence, red box), and three cp32-like P-Ps (medium confidence, blue box) from other hosts: NZ_CP115597 and NZ_CP114697 in *Borrelia miyamotoi*, and NZ_CP073149 in *Borrelia nietonii*. The cp32-like P-Ps (blue box) were only detected by tyPPing.

**Figure S12.**
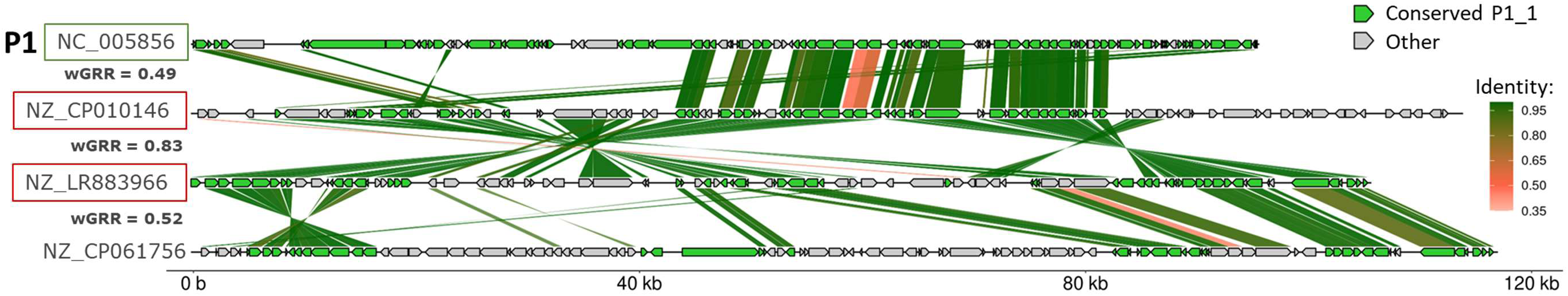
Detection of P1-like P-Ps of 03/21 by tyPPing. P1 (in green box) is compared to two elements (NZ_CP010146, NZ_LR883966) predicted by tyPPing to belong to P1_1 (medium confidence, red boxes). These elements were characterized as P1-like plasmids (18), that lack phage genes and encode either *oriT* sequences or relaxases. NZ_CP061756, also a P1-like plasmid, was not detected as P-P by tyPPing.

**Figure S13.**
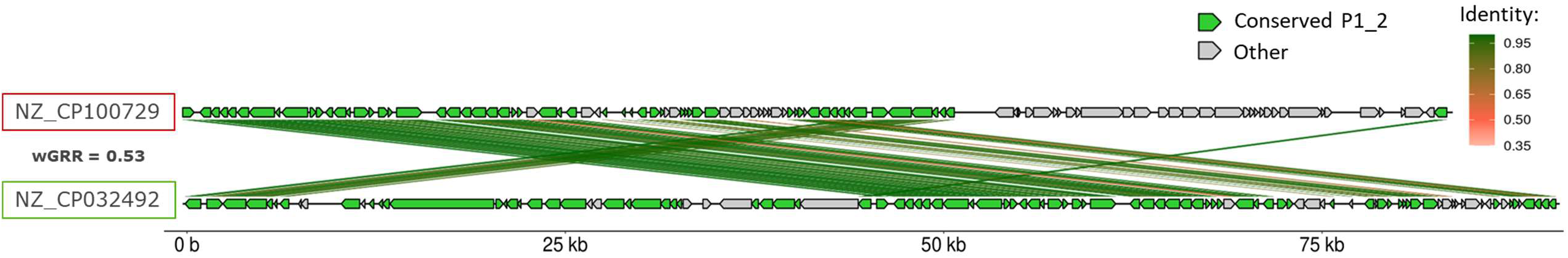
Questionable P1_2 element detected only by tyPPing in 05/23. P1_2 P-P (high confidence, green box) and an element predicted as P1_2 P-P (red box) by tyPPing. We speculate that this element is a plasmid, since it misses many P1_2 genes. Random forest models of MM-GRC assigned an average phage score of 0.496 (cutoff >0.5).

**Figure S14:**
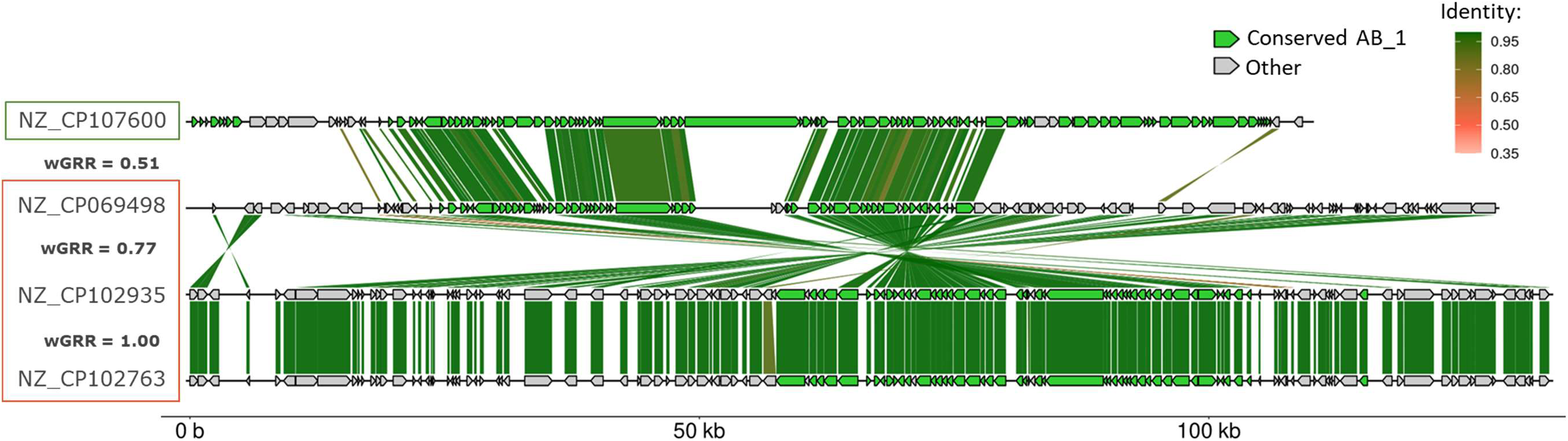
AB_1-like elements not detected by tyPPing. One high-confidence AB_1 (NZ_CP107600, green box) example and three cases (red boxes), detected by MM-GRC and not by tyPPing. These cases were not detected by tyPPing due to too few hits to AB_1 protein profiles.

**Figure S15:**
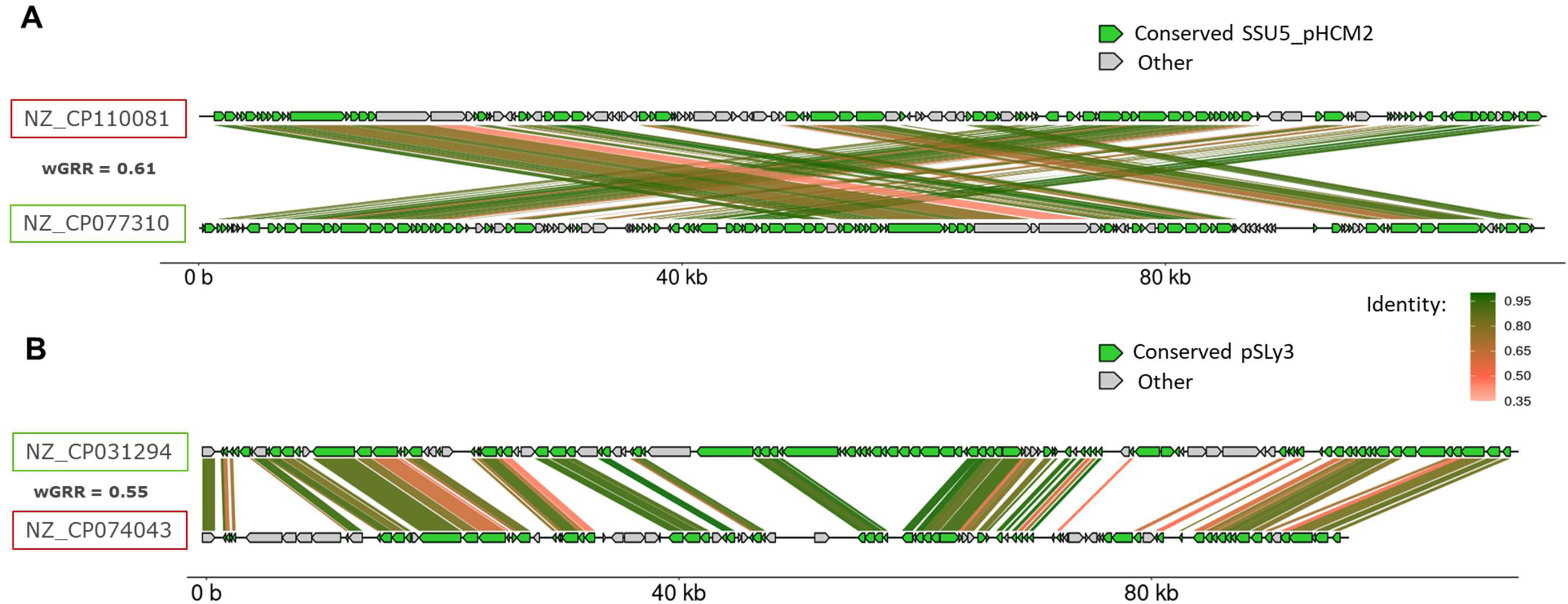
SSU5-like elements not detected by tyPPing. Two putative P-Ps, one SSU5_pHCM2-like (A, red box) and one pSLy3-like (B, red box), were missed by tyPPing A. SSU5_pHCM2-like P-P (high confidence, green box) and a similar P-P (NZ_CP110081, red box) detected in *Buttiauxella sp*. B. pSLy3-like P-P (high confidence, green box) and a related P-P (red box, NZ_CP074043) of *E. coli* strain PI6.

**Figure S16.**
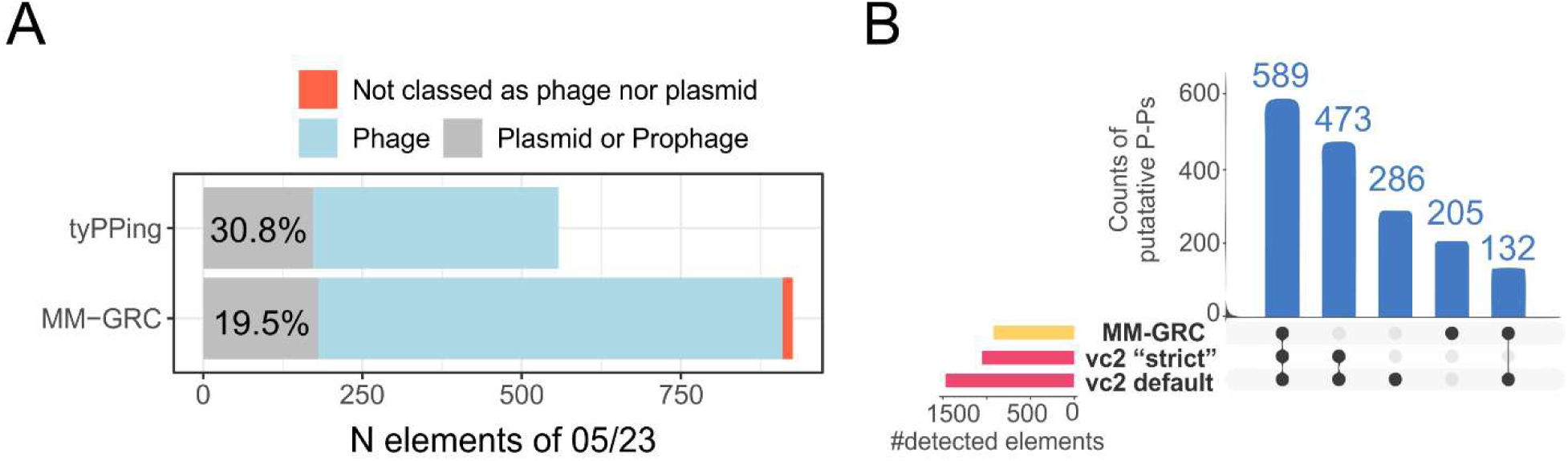
Classification of P-Ps of 05/23 using vConTACT v2 and geNomad. **(A).** Number of P-Ps predicted by tyPPing or MM-GRC classed by geNomad as phage (blue), plasmid and/or integrated prophage (grey) or not classed (red). **(B)** Counts of P-Ps predicted in 05/23 (not already present in 03/21), by using vConTACT v2 and 1416 P-Ps of 03/21 (18) as references. We compared two clustering approaches (‘strict’ and default) to predictions of MM-GRC. In ‘strict’ only cases that clearly clustered with P-Ps were considered as those. In default, cases that grouped in overlapping clusters or were categorized as Clustered/Singleton (described in the wiki of vConTACT v2 https://bitbucket.org/MAVERICLab/vcontact2), were also considered.

**Figure S17.**
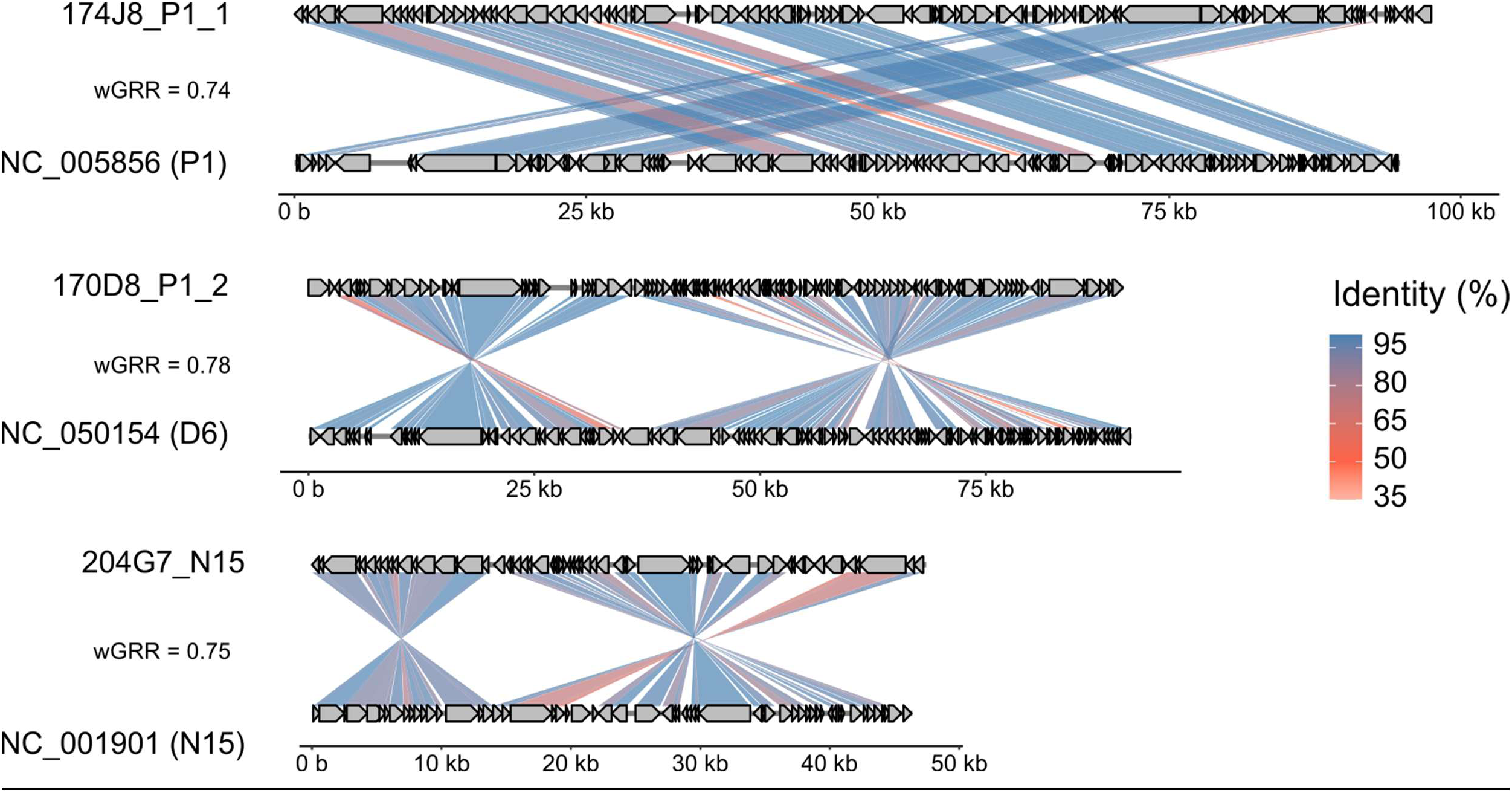
Complete assembled P-Ps of the CRE strains 174J8 (*E. coli*), 170D8 (*E. coli*) and 204G7 (*E. cloacae*). Circular, complete P-P genomes were assembled from long (PacBio) and short reads (following MMC induction, see Methods). These genomes were compared to P1 (NC005856), D6 (NC050154, P1_2), and N15 (NC001901) with gggenomes (https://github.com/thackl/gggenomes). Gene-to-gene assignments are BBHs. Pairwise protein similarity is given in %.

## References

1. Shan, X., Szabo, R.E. and Cordero, O.X. (2023) Mutation-induced infections of phage-plasmids. Nat Commun, 14, 2049.

2. Lehnherr, H., Maguin, E., Jafri, S. and Yarmolinsky, M.B. (1993) Plasmid addiction genes of bacteriophage P1: doc, which causes cell death on curing of prophage, and phd, which prevents host death when prophage is retained. J Mol Biol, 233, 414–428.

3. Dziewit, L., Jazurek, M., Drewniak, L., Baj, J. and Bartosik, D. (2007) The SXT conjugative element and linear prophage N15 encode toxin-antitoxin-stabilizing systems homologous to the tad-ata module of the *Paracoccus aminophilus* plasmid pAMI2. J Bacteriol, 189, 1983–1997.

4. Silpe, J.E. and Bassler, B.L. (2019) A Host-Produced Quorum-Sensing Autoinducer Controls a Phage Lysis-Lysogeny Decision. Cell, 176, 268–280.e13.

5. Mardanov, A.V. and Ravin, N.V. (2007) The Antirepressor Needed for Induction of Linear Plasmid-Prophage N15 Belongs to the SOS Regulon. J Bacteriol, 189, 6333–6338.

6. Yarmolinsky, M.B. (2004) Bacteriophage P1 in Retrospect and in Prospect. J Bacteriol, 186, 7025–7028.

7. Łobocka, M.B., Rose, D.J., Plunkett, G., Rusin, M., Samojedny, A., Lehnherr, H., Yarmolinsky, M.B. and Blattner, F.R. (2004) Genome of Bacteriophage P1. J Bacteriol, 186, 7032–7068.

8. Ravin, N.V. (2011) N15: The linear phage–plasmid. Plasmid, 65, 102–109.

9. Pfeifer, E., Moura de Sousa, J.A., Touchon, M. and Rocha, E.P.C. (2021) Bacteria have numerous distinctive groups of phage–plasmids with conserved phage and variable plasmid gene repertoires. Nucleic Acids Res, 49, 2655–2673.

10. Figueroa, W., Cazares, D. and Cazares, A. (2024) Phage-plasmids: missed links between mobile genetic elements. Trends Microbiol, 32, 622–623.

11. Sayid, R., van den Hurk, A.W.M., Rothschild-Rodriguez, D., Herrema, H., de Jonge, P.A. and Nobrega, F.L. (2024) Characteristics of phage-plasmids and their impact on microbial communities. Essays Biochem, 68, 583–592.

12. Szczepankowska, A.K. and Łobocka, M. (2024) Exploring the role of phage plasmids in gene transfers. Trends Genet, 40, 555–557.

13. Brisson, D., Zhou, W., Jutras, B.L., Casjens, S. and Stevenson, B. (2013) Distribution of cp32 Prophages among Lyme Disease-Causing Spirochetes and Natural Diversity of Their Lipoprotein-Encoding erp Loci. Appl Environ Microbiol, 79, 4115–4128.

14. Pfeifer, E., Bonnin, R.A. and Rocha, E.P.C. (2022) Phage-Plasmids Spread Antibiotic Resistance Genes through Infection and Lysogenic Conversion. mBio, 13, e01851–22.

15. Enault, F., Briet, A., Bouteille, L., Roux, S., Sullivan, M.B. and Petit, M.-A. (2017) Phages rarely encode antibiotic resistance genes: a cautionary tale for virome analyses. ISME J, 11, 237–247.

16. Venturini, C., Zingali, T., Wyrsch, E.R., Bowring, B., Iredell, J., Partridge, S.R. and Djordjevic, S.P. (2019) Diversity of P1 phage-like elements in multidrug resistant *Escherichia coli*. Sci Rep, 9, 18861.

17. Billard-Pomares, T., Fouteau, S., Jacquet, M.E., Roche, D., Barbe, V., Castellanos, M., Bouet, J.Y., Cruveiller, S., Médigue, C., Blanco, J., et al. (2014) Characterization of a P1-Like Bacteriophage Carrying an SHV-2 Extended-Spectrum β-Lactamase from an *Escherichia coli* Strain. Antimicrob. Agents Chemother, 58, 6550–6557.

18. Pfeifer, E. and Rocha, E.P.C. (2024) Phage-plasmids promote recombination and emergence of phages and plasmids. Nat Commun, 15, 1545.

19. Camargo, A.P., Roux, S., Schulz, F., Babinski, M., Xu, Y., Hu, B., Chain, P.S.G., Nayfach, S. and Kyrpides, N.C. (2024) Identification of mobile genetic elements with geNomad. Nat Biotechnol, 42, 1303–1312.

20. Arndt, D., Grant, J.R., Marcu, A., Sajed, T., Pon, A., Liang, Y. and Wishart, D.S. (2016) PHASTER: a better, faster version of the PHAST phage search tool. Nucleic Acids Res, 44, W16–21.

21. Redondo-Salvo, S., Bartomeus-Peñalver, R., Vielva, L., Tagg, K.A., Webb, H.E., Fernández-López, R. and de la Cruz, F. (2021) COPLA, a taxonomic classifier of plasmids. BMC Bioinform, 22, 390.

22. Carattoli, A. and Hasman, H. (2020) PlasmidFinder and In Silico pMLST: Identification and Typing of Plasmid Replicons in Whole-Genome Sequencing (WGS). Methods Mol Biol, 2075, 285–294.

23. Néron, B., Littner, E., Haudiquet, M., Perrin, A., Cury, J. and Rocha, E. (2022) IntegronFinder 2.0: Identification and Analysis of Integrons across Bacteria, with a Focus on Antibiotic Resistance in *Klebsiella*. Microorganisms, 10, 700.

24. Cury, J., Abby, S.S., Doppelt-Azeroual, O., Néron, B. and Rocha, E.P.C. (2020) Identifying Conjugative Plasmids and Integrative Conjugative Elements with CONJscan. Methods Mol Biol, 2075, 265– 283.

25. Bin Jang, H., Bolduc, B., Zablocki, O., Kuhn, J.H., Roux, S., Adriaenssens, E.M., Brister, J.R., Kropinski, A.M., Krupovic, M., Lavigne, R., et al. (2019) Taxonomic assignment of uncultivated prokaryotic virus genomes is enabled by gene-sharing networks. Nat Biotechnol, 37, 632–639.

26. Faith, D.R., Kinnersley, M., Brooks, D.M., Drecktrah, D., Hall, L.S., Luo, E., Santiago-Frangos, A., Wachter, J., Samuels, D.S. and Secor, P.R. (2024) Characterization and genomic analysis of the Lyme disease spirochete bacteriophage ϕBB-1. PLoS Pathog, 20, e1012122.

27. Peng, J. and Xu, J. (2010) Low-homology protein threading. Bioinformatics, 26, i294–i300.

28. Altschul, S.F., Madden, T.L., Schäffer, A.A., Zhang, J., Zhang, Z., Miller, W. and Lipman, D.J. (1997) Gapped BLAST and PSI-BLAST: a new generation of protein database search programs. Nucleic Acids Res, 25, 3389–3402.

29. Achtman, M., Morelli, G., Zhu, P., Wirth, T., Diehl, I., Kusecek, B., Vogler, A.J., Wagner, D.M., Allender, C.J., Easterday, W.R., et al. (2004) Microevolution and history of the plague bacillus, *Yersinia pestis*. Proc Natl Acad Sci U S A, 101, 17837–17842.

30. Terzian, P., Olo Ndela, E., Galiez, C., Lossouarn, J., Pérez Bucio, R.E., Mom, R., Toussaint, A., Petit, M.-A. and Enault, F. (2021) PHROG: families of prokaryotic virus proteins clustered using remote homology. NAR genom. bioinform, 3, lqab067.

31. Cury, J., Touchon, M. and Rocha, E.P.C. (2017) Integrative and conjugative elements and their hosts: composition, distribution and organization. Nucleic Acids Res, 45, 8943–8956.

32. Gautreau, G., Bazin, A., Gachet, M., Planel, R., Burlot, L., Dubois, M., Perrin, A., Médigue, C., Calteau, A., Cruveiller, S., et al. (2020) PPanGGOLiN: Depicting microbial diversity via a partitioned pangenome graph. PLoS Comput. Biol., 16, e1007732.

33. Steinegger, M., Meier, M., Mirdita, M., Vöhringer, H., Haunsberger, S.J. and Söding, J. (2019) HH-suite3 for fast remote homology detection and deep protein annotation. BMC Bioinformatics, 20, 473.

34. Hepner, S., Kuleshov, K., Tooming-Kunderud, A., Alig, N., Gofton, A., Casjens, S., Rollins, R.E., Dangel, A., Mourkas, E., Sheppard, S.K., et al. (2023) A high fidelity approach to assembling the complex Borrelia genome. BMC Genomics, 24, 401.

35. Dionisio, F., Zilhão, R. and Gama, J.A. (2019) Interactions between plasmids and other mobile genetic elements affect their transmission and persistence. Plasmid, 102, 29–36.

36. Garcillán-Barcia, M.P., de la Cruz, F. and Rocha, E.P.C. (2025) The extended mobility of plasmids. Nucleic Acids Res, 53, gkaf652.

37. Sternberg, N. (1990) Bacteriophage P1 cloning system for the isolation, amplification, and recovery of DNA fragments as large as 100 kilobase pairs. Proc Natl Acad Sci U S A, 87, 103–107.

38. Nayfach, S., Camargo, A.P., Schulz, F., Eloe-Fadrosh, E., Roux, S. and Kyrpides, N.C. (2021) CheckV assesses the quality and completeness of metagenome-assembled viral genomes. Nat Biotechnol, 39, 578–585.

39. Gnezda-Meijer, K., Mahne, I., Poljsak-Prijatelj, M. and Stopar, D. (2006) Host physiological status determines phage-like particle distribution in the lysate. FEMS Microbiol Ecol, 55, 136–145.

40. Murooka, Y. and Harada, T. (1979) Expansion of the host range of coliphage P1 and gene transfer from enteric bacteria to other gram-negative bacteria. Appl Environ Microbiol, 38, 754–757.

41. Wetzel, K.S., Aull, H.G., Zack, K.M., Garlena, R.A. and Hatfull, G.F. (2020) Protein-Mediated and RNA-Based Origins of Replication of Extrachromosomal Mycobacterial Prophages. mBio, 11, e00385–20.

42. Piligrimova, E.G., Kazantseva, O.A., Nikulin, N.A. and Shadrin, A.M. (2019) Bacillus Phage vB_BtS_B83 Previously Designated as a Plasmid May Represent a New Siphoviridae Genus. Viruses, 11, 624.

43. Gillis, A. and Mahillon, J. (2014) Prevalence, genetic diversity, and host range of tectiviruses among members of the *Bacillus cereus* group. Appl Environ Microbiol, 80, 4138–4152.

44. Schüler, M.A., Daniel, R. and Poehlein, A. (2024) Novel insights into phage biology of the pathogen *Clostridioides difficile* based on the active virome. Front Microbiol, 15, 1374708.

45. Lan, S.-F., Huang, C.-H., Chang, C.-H., Liao, W.-C., Lin, I.-H., Jian, W.-N., Wu, Y.-G., Chen, S.-Y. and Wong, H.-C. (2009) Characterization of a new plasmid-like prophage in a pandemic *Vibrio parahaemolyticus* O3:K6 strain. Appl Environ Microbiol, 75, 2659–2667.

46. Santoriello, F.J. and Bassler, B.L. (2025) A family of linear plasmid phages that detect a quorum-sensing autoinducer exists in multiple bacterial species. 10.1101/2025.07.30.667625.

47. Schmidtke, D.T., Hickey, A.S., Wirbel, J., Lin, J.D., Liachko, I., Sherlock, G. and Bhatt, A.S. (2025) The prototypic crAssphage is a linear phage-plasmid. Cell Host Microbe, 10.1016/j.chom.2025.07.004.

48. Kim, M.-S. and Bae, J.-W. (2018) Lysogeny is prevalent and widely distributed in the murine gut microbiota. ISME J, 12, 1127–1141.

49. Sutcliffe, S.G., Reyes, A. and Maurice, C.F. (2023) Bacteriophages playing nice: Lysogenic bacteriophage replication stable in the human gut microbiota. iScience, 26, 106007.

50. Pfeifer, E., d’Humières, C., Lamy-Besnier, Q., Oñate, F.P., Denisé, R., Dion, S., Condamine, B., Touchon, M., Ma, L., Burdet, C., et al. (2025) Antibiotic perturbation of the human gut phageome preserves its individuality and promotes blooms of virulent phages. Cell Rep, 44, 116020.

51. O’Leary, N.A., Wright, M.W., Brister, J.R., Ciufo, S., Haddad, D., McVeigh, R., Rajput, B., Robbertse, B., Smith-White, B., Ako-Adjei, D., et al. (2016) Reference sequence (RefSeq) database at NCBI: current status, taxonomic expansion, and functional annotation. Nucleic Acids Res, 44, D733– D745.

52. Moura de Sousa, J.A., Pfeifer, E., Touchon, M. and Rocha, E.P.C. (2021) Causes and Consequences of Bacteriophage Diversification via Genetic Exchanges across Lifestyles and Bacterial Taxa. Mol. Biol. Evol., 38, 2497–2512.

53. Steinegger, M. and Söding, J. (2017) MMseqs2 enables sensitive protein sequence searching for the analysis of massive data sets. Nat Biotechnol, 35, 1026–1028.

54. Eddy, S.R. (2011) Accelerated Profile HMM Searches. PLoS Comput. Biol., 7, e1002195.

55. Néron, B., Denise, R., Coluzzi, C., Touchon, M., Rocha, E.P.C. and Abby, S.S. (2023) MacSyFinder v2: Improved modelling and search engine to identify molecular systems in genomes. Peer community j, 3.

56. Eggers, C.H. and Samuels, D.S. (1999) Molecular Evidence for a New Bacteriophage of *Borrelia burgdorferi*. Journal of Bacteriology, 181, 7308–7313.

57. Katoh, K. (2002) MAFFT: a novel method for rapid multiple sequence alignment based on fast Fourier transform. Nucleic Acids Res., 30, 3059–3066.

58. Mistry, J., Chuguransky, S., Williams, L., Qureshi, M., Salazar, G.A., Sonnhammer, E.L.L., Tosatto, S.C.E., Paladin, L., Raj, S., Richardson, L.J., et al. (2021) Pfam: The protein families database in 2021. Nucleic Acids Res, 49, D412–D419.

59. Pitcher, D. g., Saunders, N. a. and Owen, R. j. (1989) Rapid extraction of bacterial genomic DNA with guanidium thiocyanate. Lett. Appl. Microbiol., 8, 151–156.

60. Kolmogorov, M., Yuan, J., Lin, Y. and Pevzner, P.A. (2019) Assembly of long, error-prone reads using repeat graphs. Nat Biotechnol, 37, 540–546.

61. Sambrook, J. and Russell, D.W. (2006) Purification of nucleic acids by extraction with phenol:chloroform. CSH Protoc, 2006, pdb.prot4455.

62. Chen, S., Zhou, Y., Chen, Y. and Gu, J. (2018) fastp: an ultra-fast all-in-one FASTQ preprocessor. Bioinformatics, 34, i884–i890.

63. Wick, R.R., Judd, L.M., Gorrie, C.L. and Holt, K.E. (2017) Unicycler: Resolving bacterial genome assemblies from short and long sequencing reads. PLoS Comput. Biol., 13, e1005595.

64. Li, H. and Durbin, R. (2009) Fast and accurate short read alignment with Burrows-Wheeler transform. Bioinformatics, 25, 1754–1760.

65. Li, H., Handsaker, B., Wysoker, A., Fennell, T., Ruan, J., Homer, N., Marth, G., Abecasis, G. and Durbin, R. (2009) The Sequence Alignment/Map format and SAMtools. Bioinformatics, 25, 2078–2079.

